# Differentiation of Large Extracellular Vesicles in Oral Fluid: Combined Protocol of Small Force Centrifugation and Pattern Analysis

**DOI:** 10.1101/2023.04.29.537961

**Authors:** Takamasa Kawano, Kohji Okamura, Hiroki Shinchi, Koji Ueda, Nomura Takeshi, Kiyotaka Shiba

**Affiliations:** Division of Protein Engineering, Cancer Institute, Japanese Foundation for Cancer Research, 3-8-31, Ariake, Koto-ku, Tokyo, 135-8550, Tokyo, Japan; Department of Oral Oncology, Oral and Maxillofacial Surgery, Tokyo Dental College, 5-11-13, Sugano, Ichikawa, Chiba, 272-8513, Japan; Department of Systems BioMedicine, National Center for Child Health and Development, 2-10-1, Okura, Setagaya-ku, Tokyo, 157-8535, Japan; Cancer Precision Medicine Center, Japanese Foundation for Cancer Research, 3-8-31 Ariake, Koto-ku, Tokyo, 135-8550, Japan

**Keywords:** EVs, large EVs, saliva, subclassification

## Abstract

Extracellular vesicles (EVs) in biofluids are highly heterogeneous entities in terms of their origins and physicochemical properties. Considering the application of EVs in diagnostic and therapeutic fields, it is of extreme importance to establish differentiating methods by which focused EV subclasses are operationally defined. Several differentiation protocols have been proposed; however, they have mainly focused on smaller types of EVs, and the heterogeneous nature of large EVs has not yet been fully explored. In this report, to classify large EVs into subgroups based on their physicochemical properties, we have developed a protocol, named EV differentiation by sedimentation patterns (ESP), in which entities in the crude large EV fraction are first moved through a density gradient of iodixanol with small centrifugation forces, and then the migration patterns of molecules through the gradients are analyzed using a non-hierarchical data clustering algorithm. Based on this method, proteins in the large EV fractions of oral fluids clustered into three groups: proteins shared with small EV cargos and enriched in immuno-related proteins (Group 1), proteins involved in energy metabolism and protein synthesis (Group 2), and proteins required for vesicle trafficking (Group 3). These observations indicate that the physiochemical properties of EVs, which are defined through low-speed gradient centrifugation, are well associated with their functions within cells. This protocol enables the detailed subclassification of EV populations that are difficult to differentiate using conventional separation methods.

## 1. INTRODUCTION

Extracellular vesicles (EVs) are defined in Minimal Information for Studies of Extracellular Vesicles (MISEV) 2018 (1) as the collective term for particles released from cells that are delineated by lipid bilayer(s) and cannot replicate by themselves. Cells have many routes to generate EVs, including multivesicular body-mediated pathways (2, 3), protrusion-originated productions (4, 5), and regulated cell death-related secretion (6). Thus, EVs contained in biofluids are quite heterogeneous populations originating from different cell types as well as distinct generation pathways (7-9).

EVs have been proposed to be involved in normal cells/organs differentiation as well as in the development and progression of various diseases (10, 11). Although small types of EVs have been gathering attention as responsible agents for these biological activities, quite a few researchers have also focused on larger EVs as central executants of biological activities (2, 12-16). Here, the sizes of EVs (small or large) can be quantitatively determined from certain measurement methods, such as nanoparticle tracking analysis, resistive pulse sensing, atomic force micrography, and transmission electron microscopy, after being differentiated by sequential centrifugation, gel filtration, field-flow fractionation, or other methods (1). However, the sizes of EVs themselves do not necessarily provide information about their origins in terms of parental cells and generation routes. For example, it is often noted that small, medium, and large sizes of EVs are generated, respectively, through multivesicular body-mediated secretion, budding from the plasma membrane, and apoptotic cell burst. However, membrane budding and apoptosis have been shown to produce small EVs (6, 17) as well as large ones, exemplifying the limitation of size-based subclassification of EVs.

In our previous work (18), we captured a whole picture of EVs contained in healthy and oral cancer patient’s oral fluids (OFs) using pentapartite differential centrifugation, in which OFs were fractionated into five sub-populations through sequential centrifugation at 300 (0.3K), 2,000 (2K), 10,000 (10K), and 160,000 (160K) *g,* with the 2K and 160K fractions enriched with large and small EVs, respectively. From the characterizations of each fraction, we found prominent alternations in the 2K fraction of patient’s OFs in terms of the amounts and the sizes of certain protein markers, which suggested that diagnostic information can be drawn not only from small but also from large EVs (18). To explore this possibility, we must first further differentiate the 2K fraction using certain defined operational manipulations, because the fractions obtained by sequential centrifugations are still very heterogeneous and can be further subclassified using other methods. For small EVs, an equilibrium density gradient centrifugation is often used for this purpose, in which EVs float from the bottom or sediment from the top through a density gradient medium until their densities agree with those of the surrounding medium (Supplementary Information 1). Although only a portion of EVs migrate to their equilibrium positions under standard conditions (19-22), this methodology has been frequently used to sort heterogeneous small EVs into subclasses with distinct densities. The key point to note for equilibrium density gradient centrifugation is that the information for the sizes of EVs is discarded at equilibrium conditions (Supplementary Information 1). In our previous observation, the large EVs contained in the 2K fraction varied in size compared to the small EVs in the 160K fraction, which suggested that we employ non-equilibrium conditions for differentiating large EVs. In this study, we propose a new protocol for differentiating large EVs, in which crude large EVs are first sedimented through density gradients at low speeds and short times, and then the migration patterns of molecules are analyzed using a non-hierarchical data clustering algorithm. We named this differentiation protocol *EV differentiation by sedimentation patterns* (ESP).

## 2. MATERIALS AND METHODS

### 2.1. Oral fluid collection

The OF samples were collected from three healthy volunteers with informed consent, as described previously (18). Briefly, 40 ml of oral fluid was expectorated into plastic tubes (2 × 20 ml) (430829, Corning, NY, USA) placed on ice. Volunteers were prohibited from eating, drinking, and brushing for 1 h prior to collection, which began at 9:00 a.m. The subjects were all males aged 28, 30, and 64 years old who did not receive any dental treatment within 48 h. The collected OFs were flash frozen in liquid nitrogen, stored at −80°C, and thawed at 4°C for use in EV isolation. Ethical approval was obtained from the Tokyo Dental College of Ethics Committee (approval number:1-16-46RIV) and the Japanese Foundation for Cancer Research (approval number: JFCR 2016-1125).

### 2.2. Differentiation of the 2K fraction

To prepare the 2K fraction, 20 ml of frozen OFs were thawed, and debris was removed by centrifugation at 300 *g* for 10 min (model 5500, Kubota, Osaka, Japan). The resultant supernatant was then centrifuged at 2,000 *g* for 10 min, and the pellets were dissolved in 5 ml of phosphate-buffered saline (PBS, 137 mM NaCl, 2.68 mM KCl, 8.10 mM Na_2_HPO_4_, 1.47 mM KH_2_PO_4_, pH 7.4) and centrifugated under the same conditions to obtain the pellet of the 2K fraction. All centrifugations were done at 4°C. In this study, we omitted the sonication step prior to 300 *g* centrifugation, which was employed in the pentapartite analyses (18), because it was only needed for the density gradient centrifugation of small EVs (21) and did not affect the properties of the 2K fraction (unpublished result). A continuous density gradient of 8–48% iodixanol (OptiPrep, 1114542, Shield PoC, Oslo, Norway) in 0.02 M HEPES ([4-(2-hydroxyethyl)-1-piperazine ethanesulfonic acid]/NaOH, pH 7.2) was prepared with a gradient mixer No. 3 (SAN4024, Sanplatec, Osaka, Japan) in a 10 ml tube (361707, Beckman Coulter). The 2K fraction was placed on the top or bottom of the media after dissolving the pellet in 500 µl of 8% or 48% iodixanol in 0.02 M HEPES, respectively. Centrifugations were performed under the various conditions indicated in the REULTS section using JXN-30 with a JS-24.38 rotor (Beckman Coulter). After centrifugation, 9 individual 1 ml fractions were collected from the top, and the density of each fraction was measured using a refractometer (RX-5000a, Atago Co. Ltd, Tokyo, Japan). Only fraction 10 (bottom) was 0.5 ml of the sample. All samples were flash frozen in liquid nitrogen, stored at −80°C, and thawed at 4°C before analysis.

### 2.3. Pentapartite fractionation

Pentapartite fractionations of OFs were performed as described by Hiraga et al. (18), with slight modifications. Briefly, 5 ml of thawed OFs were sonicated with a closed-type sonication system (UCD-200T, Biorupter, Diagenode Inc., NJ, USA) and were sequentially centrifuged at 300 *g* for 10 min (model 5500 with tube 430829, CORNING), at 2,000 *g* for 10 min (model 5500), at 10,000 *g* for 30 min (JXN-30 with JS-24.38 rotor and tube 364772, Beckman Coulter), and at 100,000 *g* for 110 min (JXN-30 with JS-24.38, 364772) to obtain the 0.3K, 2K, 10K and 100K fractions, respectively. The differences between the previous study (18) and the method described here are as follows: (i) the final pellet was obtained from 100,000 *g* for 110 min with a JS-24.38 rotor (k factor = 334) in this study instead of 160,000 *g* for 70 min with an SW32Ti rotor (k factor = 204) because of the availability of equipment; (ii) the pellets obtained from the first 300 *g*, 2,000 *g*, 10,000 *g* and 100,000 *g* centrifugation were washed once by dissolving the pellets in 5 ml (for the 0.3K and 2K fractions) or 30 ml (for the 10K and 100K fractions) of PBS, followed by centrifugation under identical conditions to reduce carry-over of preceding fractions; and (iii) 10 ml of the final supernatant (Sup) was dialyzed against 1,000 ml of Milli-Q IQ 7010 purification system (Merk, Darmstadt, Germany) water using a Slide-A-Lyzer Dialysis Cassette (66110, Thermo Fischer Scientific, MA, USA) at 4°C for 24 h by exchanging external water four times, followed by lyophilization with FREEZONE 4.5 (FZ-4.5CL, Asahi Life Science, Saitama, Japan) for 18 h to prepare the concentrated Sup fraction in 152 µl of PBS. All centrifugations were done at 4°C.

### 2.4 Transmission electron microscopy

Transmission electron microscopy (TEM) observations were outsourced to Tokai Electron Microscopy, Inc. (Nagoya, Japan), where the samples were absorbed to formvar film-coated copper grids and stained with 2% phosphate tungstic acid solution (pH 7.0) for 30 s. The grids were observed by a transmission electron microscope (JEM-1400Plus; JEOL Ltd., Tokyo, Japan) at an acceleration voltage of 100 kV. Digital images (3296 × 2472 pixels) were taken with a CCD camera (EM-14830RUBY2; JOEL Ltd, Tokyo, Japan). Particle size was calculated using the Analyze Particles tool in ImageJ (https://imagej.nih.gov/ij/).

### 2.5 Nanoparticle tracking analysis

The numbers and sizes of the particles were measured by nanoparticle tracking analysis (NTA) using a NanoSight LM10 system (Malvern Instruments, Worcestershire, UK), as previously reported (18). For pentapartite analyses, the 10 K fraction was calibrated with 150 nm diameter silica beads (24320, Polysciences, PA, USA), and 100 nm diameter beads (24041, Polysciences) were used for the 100 K fraction and supernatant. The camera level (CL) and detection threshold (DT) were set to CL 12/DT 10 and CL 14/DT 4. The samples were suspended in PBS filtered through a 0.1 µm syringe filter (6789-1301, GE Healthcare UK Ltd., Buckinghamshire, UK) and diluted to achieve the desired concentration between 2 × 10^8^ and 1 × 10^9^ particles/ml. For each sample, 30 s of capture were recorded per sample, and each measurement was performed five times independently. NTA software version 2.3 (Malvern Instruments) was used for the data analysis, and average histograms were plotted from the five measurements. Differentiated 2K fractions were measured according to the 100K fraction.

### 2.6 Mass spectrometry (MS)

We separated 600 µl each of the 10 fractions using a gradient of iodixanol (see 2.2), mixed the aliquots with 400 µl of PBS, and centrifuged them at 125,000 *g* (45,000 rpm) for 70 min (Optima Max TL with TLA-55 rotor, Beckman Coulter). The resultant sediment was dissolved in 40 µl of PBS and its protein concentration was estimated by a bicinchoninic acid (BCA) assay kit (23228, 1859078, Thermo Fisher Scientific, Waltham, MA, USA), using bovine serum albumin (23209, Thermo Fisher Scientific) as a standard. Mass spectrometric analysis of the protein samples was performed as described previously (22). Briefly, the samples were concentrated using the 2-D Clean-Up Kit (80648451, GE Healthcare, Chicago, IL), reduced in 1 × Laemmli’s sample buffer (32.9 mM Tris HCl, pH 6.8, 13.15% glycerol, 1.05% sodium dodecyl sulfate (SDS), 0.005% bromophenol blue) containing 10 mM Tris (2-carboxyethyl) phosphine for 10 min at 100°C, alkylated with 50 mM iodoacetamide at room temperature for 45 min, and subjected to SDS polyacrylamide gel electrophoresis (SDS-PAGE). The electrophoresis was stopped 2 mm from the top edge of the separation gel. After Coomassie Brilliant Blue staining, the protein bands were cut out, destained, cut into small pieces, and subjected to in-gel digestion with trypsin/Lys-C Mix (V5071, Promega, Madison, WI, USA) at 37°C for 12 h. The resulting peptides were extracted from the gel pieces and analyzed using an Orbitrap Fusion Lumos mass spectrometer (Thermo Fisher Scientific, Waltham, MA) in combination with an UltiMate 3000 RSLC nanoflow HPLC (Thermo Fisher Scientific). MS/MS spectra were searched using the SwissProt *Homo sapiens* protein sequence database with Proteome Discoverer 2.5 software (Thermo Fisher Scientific).

### 2.7 SDS-PAGE, western blot, and gel staining

For SDS-PAGE, 8 µl of sample from each fraction was incubated with 6 µl of reducing sample buffer (250 mM Tris HCl, pH 6.8, 20% sucrose, 8% SDS, 5% 2-mercaptoethanol, 0.008% bromophenol blue) and 10 µl 0.05% 2-methacryloyloxyethyl phosphorylcholine (MPC) polymer (Lipidure-BL802, NOF Corporation, Tokyo, Japan) and boiled at 90°C for 10 min (MPC polymer was added as signal enhancer). Proteins were separated on a 12.5% acrylamide gel (Extra PAGE One Precast Gel, 13074-34, Nacalai Tesque, Inc., Kyoto, Japan) using SDS running buffer (25 mM Tris, 191 mM glycine, 0.1% SDS) at a constant current of 500 V at 40 mA for 40 min. Separated proteins were then transferred onto 10 cm × 10 cm nitrocellulose membranes (IB201002, Invitrogen, CA, USA) using the iBlot dry blotting system (Invitrogen). After nonspecific binding sites were blocked by incubation in 10 ml of Blocking One (03953-95, Nacalai Tesque Corporation) for 1 h, the membrane was washed with 10 ml Tris-buffered saline-Tween [TBS-T, 10 mM Tris-HCl, 150 mM NaCl, 0.1% Tween-20 (P1379, Sigma-Aldrich)] three times for 5 min each, followed by incubation with antibody in 10 ml CanGet signal solution 1 (NKB-101, Toyobo, Osaka, Japan) for 120 min. The first antibody was probed by incubation with horseradish peroxidase-conjugated secondary antibodies in 10 ml of CanGet signal solution 2 for 60 min in the dark. All antibody incubations were performed using a gentle shaker. After each incubation step, the membranes were washed three times with 10 ml TBS-T for 5 min. Luminescence signals were obtained using an ECL system (A-8511, C-9008, Sigma-Aldrich) and visualized with a LumiVision LPR-130 (TAITEC, Saitama, Japan). For silver staining and Coomassie brilliant blue staining, Sil-Best Stain One (06865-81, Nacalai Tesque, Inc.) and InstantBlue (ISB1L, Expedeon, Cambridge, UK) were used, respectively. Antibodies used in western blotting experiments and their dilution rates were as follows: mouse anti-CD9 (SHI-EXO-M01, Cosmobio, Tokyo, Japan; 1:2000), mouse anti-CD63 (SHI-EXO-M02, Cosmobio; 1:1000), mouse anti-CD81 (SHI-EXO-M03, Cosmobio; 1:1000), rabbit anti-CD81 (66866-1-Ig, Proteintech, Rosemont, USA; 1:1000), goat anti-rabbit IgG (H + L)-HRP conjugate (170-6515, Bio-Rad, CA, USA; 1:2000), and goat anti-mouse IgG (H+L)-HRP conjugate (170-6516, Bio-Rad; 1:2000).

### 2.8 Data analysis

The data obtained from the MS results were normalized with Microsoft Excel so that the maximum value was 1 and the minimum value was 0 for each protein. The average expression levels for each of the three samples was calculated. Cluster analysis was performed using TimeSeriesKMeans (a python package based on scikit-learn (23), numpy (24), and scipy (25) published by tslearn (26)), bioinfokit (27), matplotlib (28), seaborn (29), and pandas (30). Subcellular localization and Gene Ontology analysis (31, 32) were performed using metascape (33).

### 2.9 Reporting

We have submitted all relevant data to the EV-TRACK knowledgebase (EV-TRACK ID: EV220400) (34).

## 3. RESULTS

### 3.1 Low-speed, short-time density gradient centrifugation raises the resolution of large EVs

Equilibrium density gradient centrifugation (isopycnic density gradient centrifugation) has often been used to differentiate small EVs from crude small EV fractions (2). Theoretically, in this method, each EV particle moves through a density gradient medium from the top or from the bottom of the centrifugation tube to the position where the density of the particle becomes identical to that of the medium, and the velocity of the particle converges to zero (Supplementary Information 1). Previous studies, including ours, have, however, indicated that even after 96 h centrifugation, only a limited portion of small EVs reach their equilibrium state (19-22), indicating that particles precent in the crude fraction did not behave as ideal spherical entities with distinct diameters. The observations also indicate that the non-equilibrium conditions of density gradient centrifugation (zonal density gradient centrifugation) can be used to differentiate heterogeneous EV populations based on their sizes as well as the densities of particles (Supplementary Information 1); that is, the size information of particles is not sacrificed in these conditions. This is suitable for characterizing the 2K fraction of OFs, because in our previous pentapartite analyses, TEM observations indicated that the particles contained in the 2K fraction had a wide range of sizes. For this reason, we first explored the non-equilibrium centrifugation conditions that separated EVs contained in the 2K fraction throughout the density gradient media.

According to Stokes’ law, the effect of particle size should be more prominent with decreased rotational angular velocity (Supplementary Information 1). Therefore, we first sedimented the 2K fraction from the top of an 8–48% iodixanol density gradient under three conditions: 2,000 *g* for 1 h, 2,000 *g* for 24 h, and 100,000 *g* for 1 h. After centrifugation, 10 fractions were collected from the top, and each fraction was subjected to NTA analysis and silver staining (Fig. 1B and Fig. S1). The NTA method determines the sizes of small particles based on their Brown’s motion, and it simultaneously measures the number of particles (35), leading to its wide use in characterizing small EVs. However, the method has not been optimized for larger particles, and the presence of large particles in a sample hampers particle tracking (1, 36, 37). In fact, in the analyses shown in Figure 1A, fractions 3 to 10 of 2,000 *g* for 1 h, and fractions 4 to 10 of 2,000 *g* for 24 h and 100,000 *g* for 1 h contained larger particles that hindered the analyses in these fractions (shown by bars with asterisks in Fig. 1B upper panels). By contrast, the top fractions of these samples were calculated to contain small particles by NTA, suggesting larger particles have moved toward the bottom of tubes in these centrifugation conditions. From the comparisons of NTA results obtained from the three conditions, we observed that particle dispersion increased with longer centrifugation time or higher centrifugal force, and the peak of particle concentration shifted from fraction 1 to 2 under these conditions. Silver-stained samples of fraction 5 had the strongest signals from all three conditions (Fig. 1B), and a similar result was obtained when the sample floated up from the bottom (Fig. 1C), suggesting that most but not all large EVs reached the equilibrium position (i.e., their density was 1.12–1.15 g/ml). To confirm that some EVs were in equilibrium with the medium after 2,000 *g* of centrifugation for 1 h, the presence of aquaporin 5 (AQP5) in each fraction was investigated using western blotting. AQP5 has been thought to be exclusively expressed in salivary grands in the oral cavity (38). Reflecting this, in the pentapartite assay, AQP5 was barely detected in the 0.3K fraction, the major EV source of which may be derived from detached desquamated epithelium, blood cells, and bacteria. By contrast, the presence of AQP5 in both 2K and 160K fractions was evident, suggesting that AQP5 is secreted from salivary glands into the oral space as cargos of large and small EVs. When the 2K fraction was differentiated by density gradient centrifugation under four different conditions—2,000 *g* for 1 h downward, 100,000 *g* or 1 h downward, 2,000 *g* for 1 h upward, and 100,000 *g* or 1 h upward—the strongest singles were detected in fraction 5 for all conditions, indicating that most AQP5 was carried by large EVs, whose density was 1.12 g/ml, in the 2K fraction (Fig. S2). In addition to the main signal, AQP5 was detected in the top or bottom fractions at 2,000 *g* for 1 h downward or upward, respectively. The top or bottom signal migrated downward or upward, respectively, with a prolonged centrifugation time, suggesting that portions of AQP5 were also carried by smaller EVs in the 2K fraction (Fig. S2). Assuming that the EVs that reached fraction 5 after 2,000 *g* of centrifugation for 1 h reached density equilibrium, their diameter was calculated to be approximately 5 µm. Another signal remained in fraction 1 at 2,000 *g* for 1 h downward and moved to fraction 2 at 100,000 *g* for 1 h, suggesting that the diameter of the EV in fraction 2 was approximately 250 nm. If the particles with a diameter of 250 nm have a density of 1.12 g/ml (the canonical density of small EV (2), Stokes’ law (Supplementary Information 1) indicates that such particles would remain at the fraction under the conditions of 2,000 *g* for 1 h. From these observations, we decided to use the conditions of 2,000 *g* for 1 h downward to differentiate the 2K fraction, by which heterogeneous EVs in the 2K fractions can be separated through a density gradient based mainly on their sizes.

**FIGURE 1.**
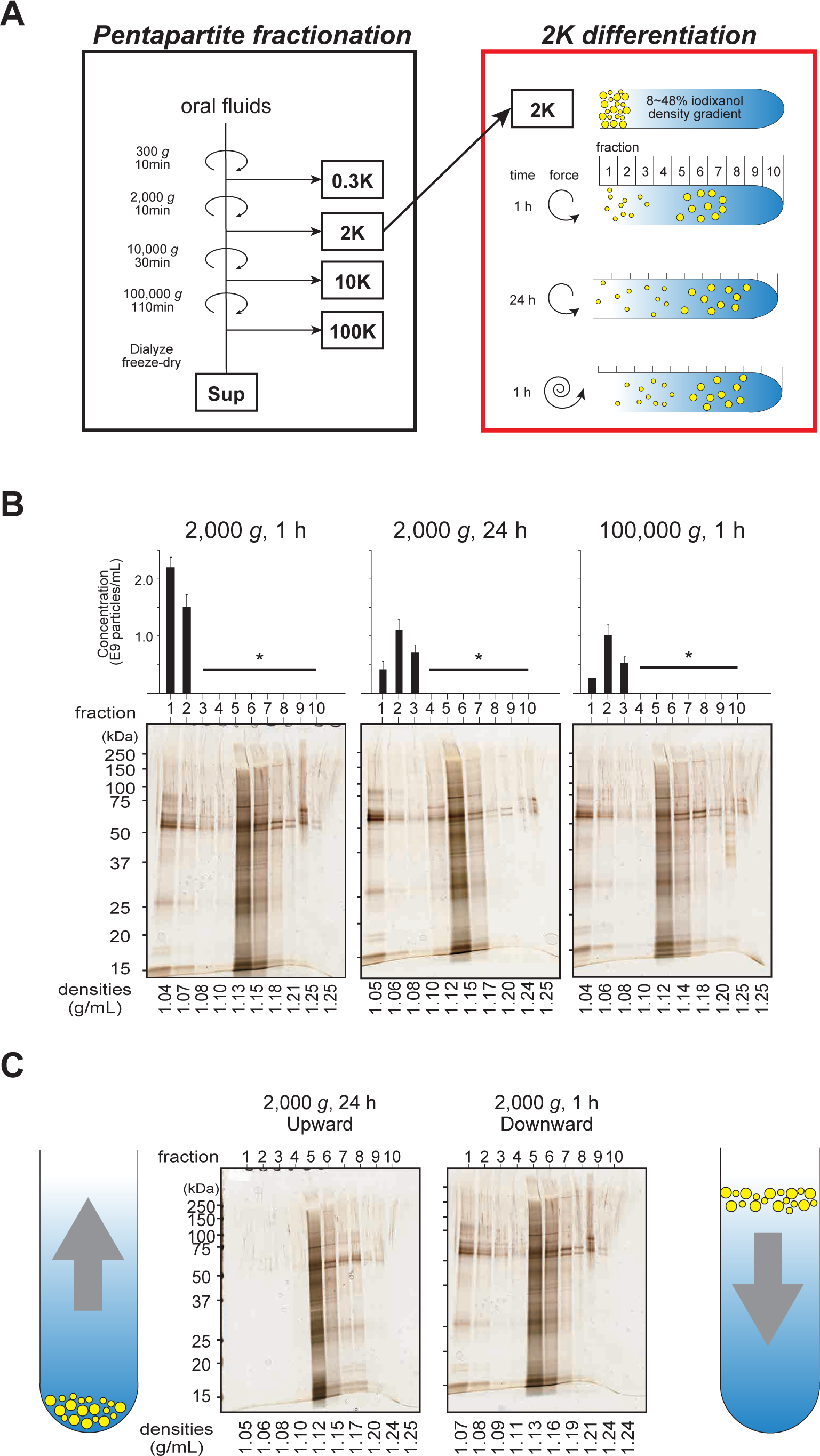
Low-speed, short-time density gradient centrifugation raises the resolution of the 2K fraction. **(A)** Schematic representation pentapartite fractionation of OFs (left) and experiments to determine the centrifugation conditions for 2K (right). The details of pentapartite fractionation were described previously; however, in this study, the previous centrifugation condition of 160,000 *g* for 70 min (k factor = 204) was changed to 100,000 *g* for 30 min (k factor = 334) because of the availability of equipment. The 2K crude fraction was further differentiated through an 8–48% iodixanol density gradient under three different conditions: 2,000 *g* for 1 h, 2,000 *g* for 24 h, and 100,000 *g* for 1 h. The length of the whorl in the right panel represents the strength of the centrifugal force. The yellow circles represent migrated particles having given sizes after centrifugations. **(B)** Concentrations of particles (upper) and silver staining of SDS-PAGE gels (bottom) of each of 10 fractions (upper) and silver staining after centrifugation under the three conditions. NTA analyses were performed five times, and the average and standard deviation are shown by black box and error bars. For fraction 1 of 100,000 *g* for 1 h, the NTA analyses were done only twice, thus the absence of an error bar. Fractions indicated by long horizontal bars with asterisks did not give concentrations because these fractions contained large particles, and these large particles hindered the measurement of NTA. Densities shown on the bottom of the gel were determined by refractometer. **(C)** Silver staining of SDS-PAGE gels for 10 fractions prepared under two different conditions. Under one condition (left), samples were floated from the bottom through gradient (upward) and under another condition (right), samples were sedimented from the top (downward).

### 3.2 Do small EVs in the 2K fraction belong to the same class of small EVs in the 160K fraction?

As shown above, in the 2K fraction, AQP5 was carried by at least two distinct types of EV; one seems to be small and the other is large. This observation suggests the possibility that the large EVs containing AQP5 represent the secreted multivesicular bodies (MVBs), in which intralumenal vesicles (small EVs) carry AQP5. The small AQP5^+^ EVs in 2K could have been those somehow released or leaked from the MVBs during the experimental procedures. Indeed, our study (18) and others’ (39-41) observations confirmed the presence of secreted MVB-like vesicles in bodily fluids.

To further explore this issue, we observed the particles contained in fractions 1, 5, and 7 using transmission electron microscopy (TEM) (Fig. 2A), and their particle sizes were measured (Fig. 2B and C). From fraction 1, small particles were mainly detected, whereas in fractions 5 and 7, much larger vesicles were detected. Unexpectedly, in fractions 5 and 7, not only large particles but also small particles were present (Fig. 2A and C). The presence of small EVs in fractions 5 and 7 supports the possibility that these small EVs rapidly migrated to these fractions by associating with large EVs, or the small EVs might have assembled into large particles under the conditions of centrifugation (42).

**FIGURE 2.**
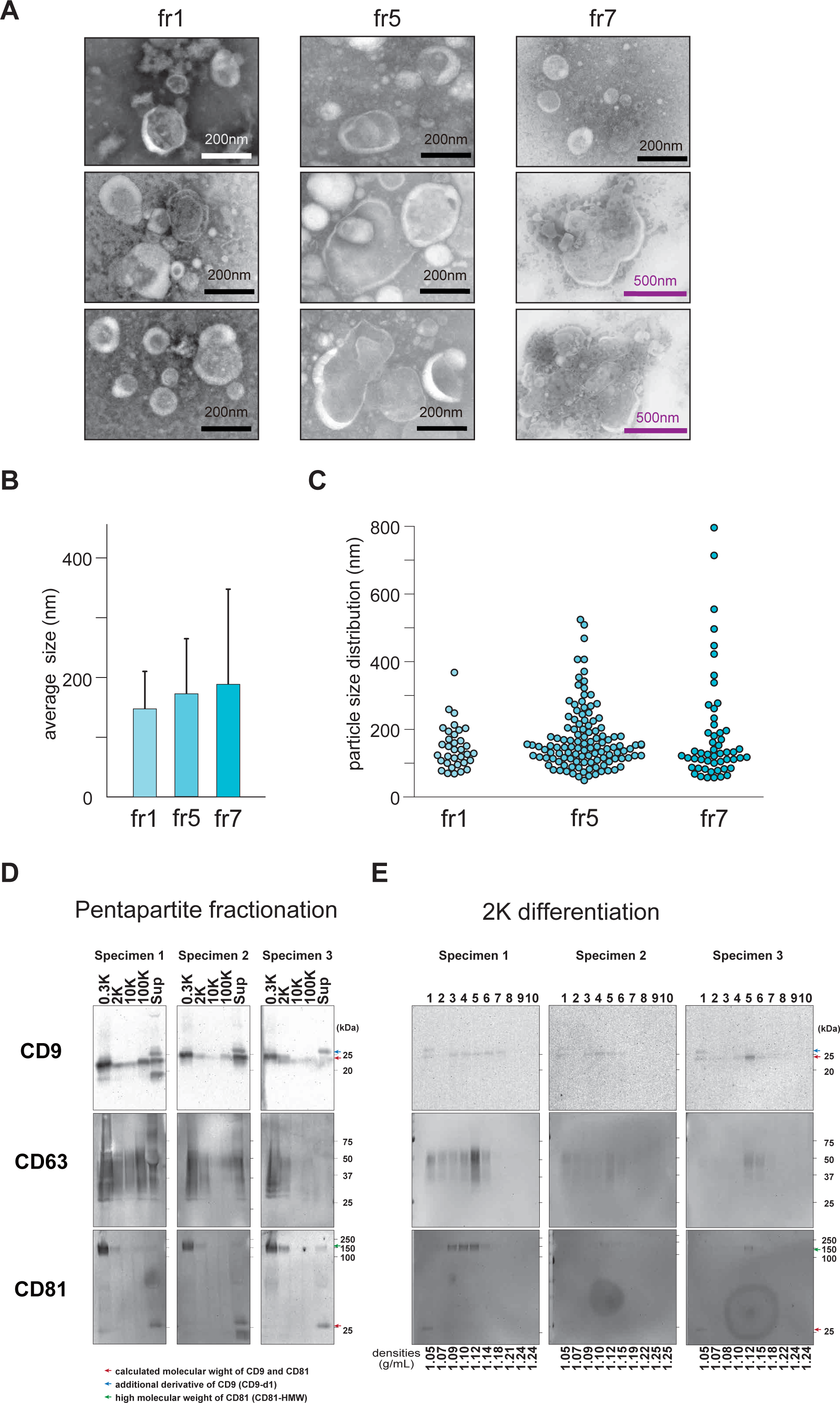
TEM images of fractions 1, 5, and 7 and distribution of CD9, CD9-d, CD63, CD81, and CD81-HMW in pentapartite fractionation and non-equilibrium density gradient centrifugation followed by western blotting. **(A)** TEM images of fractions 1, 5, and 7 after centrifugation of the 2K fraction through an 8–48% iodixanol density gradient at 2,000 *g* for 1 h. The samples were stained with 2% phosphate tungstic acid. More than 20 images were taken for each fraction, and 3 representative images were shown. Note that the scale bar is 200 nm, but the scale of the middle and right columns of fraction 7 is 500 nm. **(B)** Mean particle size (nm) of each fraction calculated using ImageJ. Error bars indicate standard deviation. There was no statistically significant difference. **(C)** Graph showing the particle size distribution in fractions 1, 5 and 7. All fractions contained particles whose sizes were within the rages of that of small EVs. **(D)** Western blots of CD9, CD63, and CD81 for the pentapartite fractions from OFs. **(E)** The 2K fractions were further differentiated by non-equilibrium density gradient centrifugation and proven with the same antibodies as (E). Numbers on the right of (D) and (E) show the molecular weights of markers. Red, blue, and green arrows indicate the calculated molecular weights of CD9 and CD81, an additional derivative of CD9 (CD9-d1), and the high molecular weight of CD81 (CD81-HMW), respectively. The origin of the CD81-HMW signal is discussed in Fig. S5.

CD9, CD63, and CD81 are other EV markers that have been detected in large EV fractions as well as small EVs (10, 13, 18, 43). In this work, we confirmed the wide distributions of these molecules with modified (see Materials and Methods) pentapartite analyses of OFs—that is, they were present on both a large EV fraction (2K) and a small EV fraction (100K) (Fig. 2D). Further, we observed the following derivatives: (i) in the Sup fraction, CD9 has given the additional derivative (CD9-d1) whose apparent molecular weight was higher than the ones in other fractions, and (ii) anti-CD81 antibody detected high molecular weight (approximately 150KDa; the origin of this high molecular weight signal was discussed in Fig. S6) signal (CD81-HMW) in 0.3K, 2K, and Sup fractions, and the signals of the calculated molecular weight were most abundant in the Sup fractions). The 2K fractions of these three specimens were further differentiated by low-speed and short-time density gradient centrifugation, as described above (Fig. 2E). Interestingly, CD9-d1, which was only detected in the Sup fraction in pentapartite analyses (Fig. 2D), was detected after this centrifugation, and this CD9-d1 was only detected in fraction 1 (Fig. 2E). The CD9 signal with calculated molecular weights was distributed with biphasic signals, whose peaks were fractions 1 and 5. CD63 exhibited monophasic distribution peaking in fraction 5. In the case of CD81, the signal with the calculated molecular weight was detected only from fraction 1, whereas CD81-HMW was detected from fractions 3 to 5. The fact that different forms of these molecules were detected from fractions 1 and 5 contradicts the simple scenario that a small EV in fraction 1 originated from the MVBs in fraction 5, as considered above. Combined with the data for AQP5 (Fig. S2), these observations indicate that distinct classes of EVs are involved in carrying AQP5, CD9, and CD81 in OFs. In the case of CD9 and CD81, different forms of these molecules (CD9 and CD9-d1, as well as CD81 and CD81-HMW) are associated with distinct EVs, highlighting the complexity of EV differentiations.

### 3.3 Classification of proteins in 2K based on their migration patterns

As shown above, the migration patterns of AQP5, CD9, and CD81 were biphasic, whereas that of CD63 was monophasic after low-speed and short-time density gradient centrifugation of the 2K fraction. We have observed various migration patterns in the proteins examined, and some are shown in Figure S3. VMP1, which is involved in autophagosome formation (44), was mostly detected in fractions 3–5. This molecule has two forms, approximately 40K and 37K, and in one specimen, the 40K form was enriched in fraction 1, resembling the case of CD9. Further, ATP5A (a subunit of the mitochondrial ATP synthase) showed a monophasic distribution and was mainly detected in fraction 6 (Fig. S3). We thought the different migration patterns through density gradient centrifugation at low speed and for a short time could offer information on their origins and decided to comprehensively examine the distributions of proteins among the 10 fractions using MS, and a total of 3,973 proteins that have been identified by MS from three OF specimens. First, we analyzed the obtained MS data using a standard hierarchical clustering method (29), which did not resolve proteins into prominent sub-groups (Fig. S4A). Next, we employed a hierarchical clustering method, TimeSeriesKMeans, which allows clustering on time-series data, such as electrocardiograms, and has been widely used for pattern analyses (26). For TimeSeriesKMeans analyses, the number of clusters had to be first determined, which we calculated to be three from our Silhouette analysis (Fig. 3A) (45). Importantly, when we compared clustering numbers 2, 3, and 4, the three groups showed the most distinct patterns (Fig. 3B, S4B).

**FIGURE 3.**
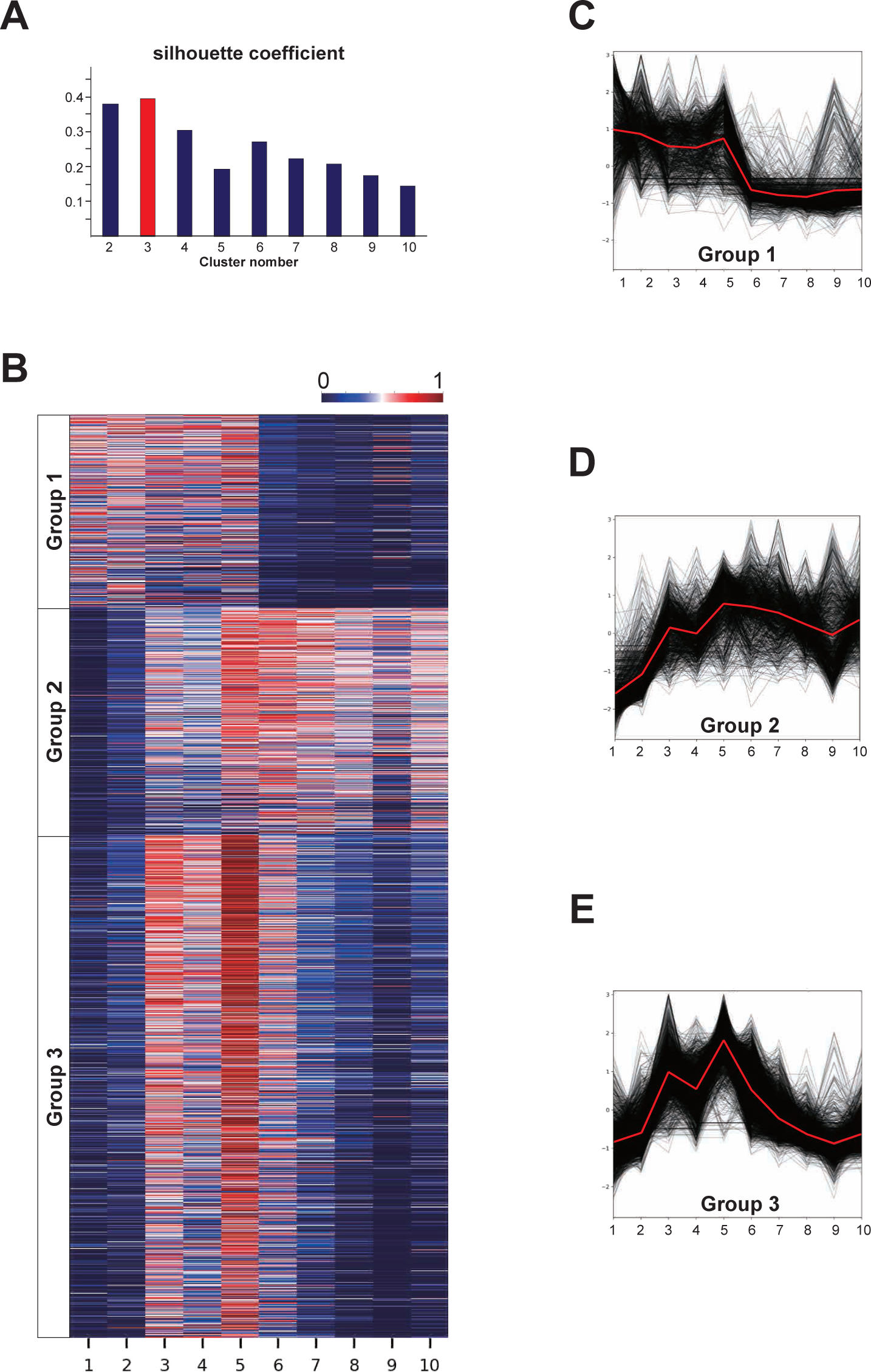
Proteins in the 2K fraction cluster into three groups based on TimeSeriesKMeans algorithm. **(A)** Silhouette analysis estimated the best number of clusters. 3. The number at the axis of the ordinate represents silhouette coefficient of the Silhouette analysis. **(B)** Heatmap of proteins clustered into three by TimeSeriesKMeans. Each cluster is named Group 1, 2, and 3. Each colored bar (total 3,973) represents a single protein, and each row represents a fraction obtained from a non-equilibrium density gradient centrifugation. Expression per protein was normalized to 0 to 1. **(C–E)** Alternative representation of expression distribution among 10 fractions. Each black line represents individual proteins, which are shown for each group. The red lines represent the centroid of the group. **(F)** Subcellular localization of proteins included in each group, which were annotated with metascape, and only those with “enhanced” and “supported” levels of evidence were used. Yellow: endomembrane system; blue: cytoplasm; green: mitochondria; red: cell nuclei; gray: not identified. The number of proteins belonging to Groups 1, 2, and 3 are 833, 978 and 2,162, respectively.

The three clusters calculated using TimeSeriesKMeans were “Group 1,” “Group 2,” and “Group 3.” each of which consisted of 833, 978 and 2,162 proteins, respectively. Proteins belonging to Group 1 were mostly present in fractions 1–5; that is, they had a low sedimentation rate (Fig. 3C, supplementary table 1). The migration pattern of the proteins in Group 2 was complementary to that of Group 1 (Fig. 3D, supplementary table 1). The proteins classified as Group 3 had bipolar peaks in fractions 3 and 5 (Fig. 3E, supplementary table 1). We name this combination of a low speed, short time gradient centrifugation, and a non-hierarchical data clustering analysis *EV differentiation by sedimentation patterns* (ESP).

### 3.4 ESP revealed three distinct functional groups of EVs in the 2K fraction

Proteins classified into three groups by ESP were annotated their subcellular localizations using metascape (33), which confirmed the segregated properties of the three groups in terms of protein localizations. Group 2 was relatively enriched with the proteins located in the mitochondria, and group 1 contained fewer cytoplasmic and nuclear-localized proteins than others (Fig. 4A). Next, using gene ontology analysis (31, 32), the proteins of the three groups were classified from three perspectives: biological processes, cellular components, and molecular functions (Supplementary table 2). The top 20 Gene Ontology terms from these analyses were evaluated for their overlapping among the three groups (Fig. 4B), which also indicated that the three groups were well segregated in terms of their cellular component, biological process, and molecular function (Fig. 4B).

**FIGURE 4.**
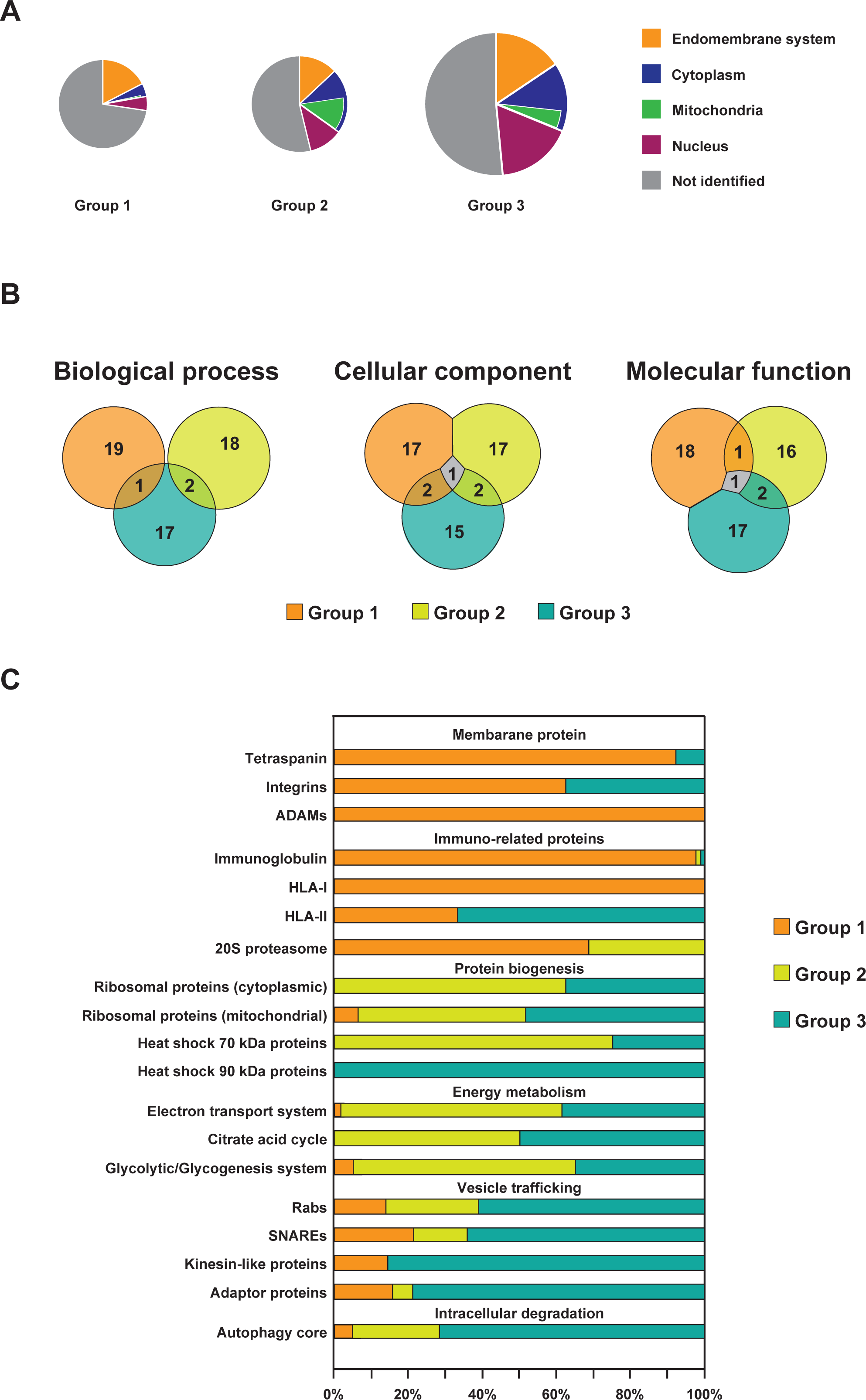
Different biological functions are related to abundant proteins in Groups 1, 2, and 3. **(A)** Venn diagrams of results obtained from Gene Ontology enrichment analysis using top 20 protein from each group (Supplementary table 2). Metascape was used to annotate cellular components, biological processes, and molecular functions, and the overlaps between the groups are shown. **(B)** Distribution protein families among three groups are shown by bar graphs, which are grouped by protein families, that is, membrane protein, protein biogenesis, vesicle trafficking, intracellular degradation, immune, and energy metabolism. Orange: Group 1, yellow: Group 2, blue: Group 3.

Next, we focused on a specific protein family, and we investigated the distribution of a group of proteins belonging to a certain protein family across the three groups (Fig. 4C). Tetraspanins, including CD9, 63, and 81, which are often referred to as small EV markers but indeed are present in large EV fractions (13, 18), were enriched in Group 1 (**12**/0/1, representing the number of proteins belonging to Group 1/2/3, respectively. The most abundant group is shown in bold letters. The same applies hereinafter. Complete data are shown in Supplementary table 3). Integrins and a disintegrin and metalloproteinases (ADAMs) are proteins regarded as small EV markers in some reports, while others reported them to be present in both small and large EVs (13, 46). These proteins also characterize Group 1 (**5**/0/3 and **3**/0/0, respectively), other examples of which include those involved in immuno-related systems: immunoglobulins and HLA-Is were almost exclusively classified into Group 1 (**85**/1/1, and **3**/0/0, respectively). The 20S proteasome proteins, which are involved in processing antigenic peptides for the presentation by HLA-I (47), were also enriched in Group 1 (**11**/5/0). In Groups 2 and 3, the proteins involved in protein biogenesis and energy metabolism were abundant, including cytoplasmic ribosomal proteins (2/**62**/44), mitochondrial ribosomal proteins (0/**48**/29), the heat shock 70 kDa protein family (1/**6/**2), electron transport system (0/**34**/26), citrate acid cycle (0/**11**/**8**), and glycolytic system (2/**10**/4). The proteins involved in protein biogenesis and energy metabolism are somewhat more prevalent in Group 2, but a considerable number of family proteins is also present in Group 3. Therefore, EVs containing these proteins may migrate during low-speed, short-time density gradient centrifugation with the patterns of Groups 2 and 3.

The fact that proteins engaging in protein biogenesis and energy metabolism were biased toward both Groups 2 and 3 might suggest that Groups 1 and 2 are not well segregated by migration pattern-based analysis. However, when focusing on other protein families, it becomes evident that Groups 2 and 3 are indeed characterized by proteins with distinct functionalities: Protein families related to vesicle trafficking, such as Rabs (5/9/**22**), SNAREs (3/2**/9**), Kinesin-like proteins (1/0/**6**), and adaptor proteins (3/1/**15**), were all enriched in Group 3. The proteins categorized into the core autophagy system (48) were also enriched in Group 3 (4/20/**61**). It should be noted that HSP90, which is known to stabilize Beclin-1 (49), a key regulator of autophagy, was enriched in Group-3 (0/0**/4**).

Overall, Groups 1, 2, and 3, which were classified based on their migration patterns following low-speed, short-time density gradient centrifugation, are characterized by the following properties: Group 1 was enriched with immuno-related proteins, and it shares EV marker proteins (tetraspanins and integrins) with small EVs, and Group 2 was characterized by protein biosynthesis and energy metabolism proteins. While Group 3 exhibits some of these proteins, its prominent characteristic was the enrichment of vesicle trafficking-related and autophagy-related proteins.

## 4. DISCUSSION

The heterogeneity of EVs stem from (1) the differences in the parent cells from which they are derived, (2) the differences in their generation pathways, and (3) the stochastic loading of cargos on the EVs. In the early phase of EV research in this decade, small EVs have gathered attention, often mentioned as generated through multivesicular bodies and uncritically named exosomes (50). Thus, only the endosome-dependent pathway has been studied in relation to EV-generation pathways. However, recent studies have revealed that cells have a tremendous number of different systems that can produce EVs (1, 51). A comprehensive understanding of these pathways has not yet been achieved; however, some of the identified generation routes can be broadly categorized into three classes: (i) endosomal organelle-dependent pathways (2, 3), (ii) protruded structure-derived pathways (4, 5), and (iii) regulated cell death pathways (6). These pathways produce EVs that have a wide range of sizes. For example, one of the regulated cell death pathways, apoptosis, is known to produce very large EVs, but the pathway also generates small EVs (52). Therefore, the size of EVs cannot sufficiently inform the routes responsible for their generation. Thus, the size fractionation of EVs is not solely sufficient for differentiating complex EV populations at high resolution. To solve this issue, equilibrium density gradient centrifugation has often been used to further differentiate crude small EV populations into their subclasses, depending on their densities (2). In this methodology, size information of EVs is abandoned in the equilibrium state, which is not problematic in the case of small EVs because they have relatively uniform sizes. However, previous studies, including ours, have shown that the fraction that contains large EVs also include small EVs. Given that small EVs cannot be sedimented under typically centrifugation conditions (for instance, 2,000 *g* for 10 min), they are somewhat associated with larger entities (or artificially generated during the EV preparation process). The coexistence of small EVs and large EVs in large EV fractions encouraged us to separate these EVs under the non-equilibrium conditions of density gradient centrifugation so that the size information would contribute to the differentiation.

When the 2K fraction of OFs was sedimented through a gradient of iodixanol under the conditions of 2,000 *g* for 1 h, small EVs remained in the top fractions, and large EVs migrated to the middle position, enabling separation of the size different EVs. Interestingly, TEM observations indicated that the fractions of large EVs (fast migrating fractions) also contained large numbers of small EVs. This fact supports the possibility that small EVs became bound to large EVs and behaved as larger particles during centrifugation. Some portions were detached from the complex and acted as free small particles. However, as shown in this study, the origins of the small EV in the large EV fraction were much more complex. For instance, western blotting results showed that CD9 and CD81 are present in different forms in terms of their apparent molecular weight (Fig. 2D). CD9 gave the additional signal that had a slightly higher molecular weight; CD9-d1 and anti-CD81 antibody detected the high molecular weight CD81-HMW. Previous studies have reported that CD9 has its palmitoylated form (53, 54), which may correspond to CD9-d1. The CD81-HMW was only detected in OF but not from plasma samples or the conditioned medium of cell lines (unpublished data) and was detected in some OF specimens. Both CD81 and CD81-HMW were recognized by two different antibodies, SHI-EXO-M03 and 66866-1-Ig (whose epitopes are residues 35-54 and 114-199 of CD81, respectively) (Fig. S6A), excluding the possibility that the signal was antibody-specific background noise. Given that CD81 has been known to act as a chaperone for CD19 (55), we evaluated the possibility that CD81-HMW contains CD9 as a component; however, wester blotting with anti-CD19 has shown that this is not the case (Fig. S6B). Glycosylation is one possibility that could have increased the molecular weight of CD81, although no study has reported the existence of glycosylated forms of CD81. Treatments of the 2K fraction with PNGaseF or O-glycosidase, which cleaves N-linked glycans or O-linked glycans from glycoproteins, respectively, did not decrease the apparent molecular weight of CD81-HMW (Fig. S6C–E), dismissing the possibility that heavy glycosylation of CD81 increased the molecular weight. Quite a few membranous proteins have been reported to migrate anomalously in SDS-PAGE. In some cases, heat treatment of a sample that is included in a standard SDS-PAGE protocol has an adverse effect on the results (56). To investigate the effect of heat treatment, the proteins in 2K were separated by diagonal 2D SDS-PAGE, which was used to detect the effects of thermal denaturation (Fig. S6F– K) (57). Diagonal migration of proteins indicated that the CD81-HMW was not an artifact generated during sample preparation (for more detailed discussion, see legend of Fig. S5). Although the origin of CD81-HMW is not explained by previously known mechanisms, two molecular forms of CD81 and CD9 provide valuable clues as to the nature of the small EVs in the large EV fraction. As shown in the Figure 2E, when the 2K fraction was developed by a non-equilibrium density gradient, two forms of these proteins were segregated into different fractions: CD9-d1 and CD81 remained in fraction 1, and CD9 and CD81-HMW moved to fractions containing large EVs. Since the large EV fraction contained many small EVs (Fig. 2C), the segregation of two forms of CD9 and CD81 ruled out the simple scenario in which small EVs become bound to the surfaces of large EVs and some small EVs behaved as larger particles and others acted as free particles after detaching from large EV in the centrifugation. With the current observations, we could not rule out the improbable possibility that CD9 and CD81-HMW in fractions 3 to 5 stemmed only from large EVs but not from small EVs. Considering the fact that the small EVs in the small EV crude fraction (160K) have been further differentiated into several subclasses, distinct types of small EVs exist in OFs not only in the small EV fraction but also in the large EV fraction.

Non-equilibrium density gradient centrifugation at 2,000 *g* for 1 h separates EVs based on their sizes and densities. As discussed above, a single EV-generation pathway could produce various sizes of EVs, and most probably, with various densities. However, we were interested in the possibility that certain EV-generation pathways might have characteristic distributions of sizes or densities of EVs (for instance, one pathway always produces both small and large EVs, the other generates a large and high density of EVs, and so on). Therefore, we intended to elucidate the characteristic patterns of molecular distributions in non-equilibrium density gradient centrifugation. We named this combination of biochemical separation and data analysis *EV differentiation by sedimentation patterns* (ESP). After separation of entities in the 2K fraction into 10 fractions with a low speed and short time gradient centrifugation, the proteins contained in each fraction were identified by MS. Standard clustering analyses of MS data did not detect prominent groupings of the segregated proteins (Fig. S4). However, when the data were analyzed by TimeSeriesKMeans, three well-separated groups were revealed (Fig. 3B). TimeSeriesKMeans is an algorithm that has been developed for clustering high-dimensional data and possesses high performance in searching patterns from data (26). The proteins that were sorted into Group 1 were abundant in fractions 1 to 5, which included many membrane proteins, such as CD9, CD63, CD81, and ADAMs. Proteins involved in protein biosynthesis and energy metabolism were enriched in Group 2, and they were moved to fractions 5–10. In Group 3, the proteins involved in protein biosynthesis and energy metabolism were also present, but they are distinguished from Group 2 via protein enrichment related to vesicle trafficking and autophagy. The migration pattern of Group 3 demonstrated bipolar peaks in fractions 3 and 5 (Fig. 3C–E).

Yamamoto et al. performed proteomic analysis of OFs’ fractions that contained medium and small EVs (22). Comparison of proteome results from these two data provides information about overlapping and independency between the crude large EV fraction and the crude medium and small EV fraction (Fig. S7). Among these fractions, 961 out of 3,973 (this study) and 1,429 (Yamamoto et al. (22)) proteins overlapped, among which 421, 210, and 330 proteins belonged to Groups 1, 2, and 3 were identified in this study (Fig. S6). In other words, half of the proteins that were sorted into Group 1 were also present in middle to small EVs, while the majority of proteins classified into Groups 2 and 3 were specific to the 2K fraction. In previous studies, proteomic analysis and western blotting showed that mitofilin was enriched in large EVs compared to small EVs (13, 17), and ADAM10 was enriched in low-density small EVs (17). In this study, mitofilin was distributed in Group 2, which contains many proteins specific to the 2K fraction, and ADAM10 belonged to Group 1, which contains many proteins that have also been used as small EV markers (13). Mitofilin was not detected in the experiment by Yamamoto et al. (22). ACTN4 (13), and ANXA1 (46) were reported as large EV markers, and they were classified into Group 2. From these data, Group 2 and Group 3 proteins have characteristic features of the large EV fraction. When these proteins were assigned their annotated characters, we found functional relationships between Group 1 and immuno-related and membrane proteins, Group 2 and energy metabolism and translation, and Group 3 and vesicle trafficking. These correlations may reflect the specific generation routes of EVs, as discussed above.

In the pentapartite fractionation by Hiraga et al. (18), the Sup fraction, the final supernatant, was for western blotting without concentration, making it difficult to directly compare the abundance of proteins between the EV fractions and supernatant. In this study, we modified the pentapartite fractionation protocol to concentrate Sup so that a more intuitive visual comparison was possible. Interestingly, the concentrated Sup demonstrated a comparable or even stronger signal than the 100K fraction for CD9, CD63, and CD81 (Fig. 2D). Recent studies have revealed that exomeres (58) and supermeres (59) also carry these molecules, which are tiny particles recovered in supernatant fractions in standard EV preparation protocol and, based on MISEV definition of EV, are non-EV particles because they do not have typical lipid bilayers found in vesicles. Thus, CD9, CD63, and CD81, which used to be regarded as small EV markers, are widely distributed among various particles in bodily fluids. However, as shown in this work and in previous work, some of their derivatives (such as CD9-d1 or CD81-HMW) show varied distributions, which may contribute to the development of EV-based diagnosis (60).

## ACKNOWKEDGEMENTS

This work was supported by JSPS KAKENHI Grant Number 20H03538. The authors thank C. Hiraga (Tokyo Dental College) and T. Minamisawa (Japanese Foundation for Cancer Research) for their helpful discussions. The authors acknowledge NOF Corporation for providing Lipidure-BL802.

## AUTHORS CONTRIBUTIONS

Takamasa Kawano contributed to the experimental performance, data acquisition and analysis, and manuscript writing, and Kohji Okamura contributed to data analysis. Hiroki Shinchi and Koji Ueda contributed to data acquisition and the analysis of MS, while Nomura Takeshi contributed to the study concept and Kiyotaka Shiba contributed to the study concept and design, data interpretation, and manuscript revision. All authors have read and approved the current version of the manuscript.

### Table of Contents (TOC)

Non-equilibrium centrifugation through the density gradient at low speed and short time followed by analyzing the distribution patterns or proteins (EV differentiation by sedimentation patterns, ESP) segregated the proteins included in the large extracellular vesicles fraction of oral fluids into Group 1, Group 2, and Group 3. Each group is enriched with proteins of specific functions, cellular location, and family, indicating that the EVs generated from different pathways and processes have distinct characteristic physical properties.

## REFERENCES

1. C. Théry et al., Minimal information for studies of extracellular vesicles 2018 (MISEV2018): a position statement of the International Society for Extracellular Vesicles and update of the MISEV2014 guidelines. J Extracell Vesicles 7, 1535750 (2018).

2. G. Raposo et al., B lymphocytes secrete antigen-presenting vesicles. J Exp Med 183, 1161–1172 (1996).

3. R. M. Johnstone, Revisiting the road to the discovery of exosomes. Blood Cells Mol Dis 34, 214–219 (2005).

4. B. H. Sung, C. A. Parent, A. M. Weaver, Extracellular vesicles: critical players during cell migration. Dev Cell 56, 1861–1874 (2021).

5. K. Rilla, Diverse plasma membrane protrusions act as platforms for extracellular vesicle shedding. J Extracell Vesicles 10, e12148 (2021).

6. I. K. H. Poon et al., Moving beyond size and phosphatidylserine exposure: evidence for a diversity of apoptotic cell-derived extracellular vesicles *in vitro*. J Extracell Vesicles 8, 1608786 (2019).

7. M. Colombo, G. Raposo, C. Théry, Biogenesis, secretion, and intercellular interactions of exosomes and other extracellular vesicles. Annu Rev Cell Dev Biol 30, 255–289 (2014).

8. G. van Niel, G. D’Angelo, G. Raposo, Shedding light on the cell biology of extracellular vesicles. Nat Rev Mol Cell Biol 19, 213–228 (2018).

9. E. I. Buzas, The roles of extracellular vesicles in the immune system. Nat Rev Immunol 23, 236–250 (2023).

10. A. Lischnig, M. Bergqvist, T. Ochiya, C. Lasser, Quantitative proteomics identifies proteins enriched in large and small extracellular vesicles. Mol Cell Proteomics 21, 100273 (2022).

11. C. Salomon et al., Extracellular vesicles and their emerging roles as cellular messengers in endocrinology: An Endocrine Society Scientific Statement. Endocr Rev 43, 441–468 (2022).

12. H. C. Anderson, Vesicles associated with calcification in the matrix of epiphyseal cartilage. J Cell Biol 41, 59–72 (1969).

13. J. Kowal et al., Proteomic comparison defines novel markers to characterize heterogeneous populations of extracellular vesicle subtypes. Proc Natl Acad Sci U S A 113, E968–977 (2016).

14. I. Melentijevic et al., *C. elegans* neurons jettison protein aggregates and mitochondria under neurotoxic stress. Nature 542, 367–371 (2017).

15. S. Kilinc et al., Oncogene-regulated release of extracellular vesicles. Dev Cell 56, 1989–2006 e1986 (2021).

16. A. D. Weems et al., Blebs promote cell survival by assembling oncogenic signalling hubs. Nature 615, 517–525 (2023).

17. R. Crescitelli et al., Subpopulations of extracellular vesicles from human metastatic melanoma tissue identified by quantitative proteomics after optimized isolation. J Extracell Vesicles 9, 1722433 (2020).

18. C. Hiraga et al., Pentapartite fractionation of particles in oral fluids by differential centrifugation. Sci Rep 11, 3326 (2021).

19. M. Aalberts et al., Identification of distinct populations of prostasomes that differentially express prostate stem cell antigen, annexin A1, and GLIPR2 in humans. Biol Reprod 86, 82 (2012).

20. J. Palma et al., MicroRNAs are exported from malignant cells in customized particles. Nucleic Acids Res 40, 9125–9138 (2012).

21. K. Iwai, T. Minamisawa, K. Suga, Y. Yajima, K. Shiba, Isolation of human salivary extracellular vesicles by iodixanol density gradient ultracentrifugation and their characterizations. J Extracell Vesicles 5, 30829 (2016).

22. S. Yamamoto et al., Specimen-specific drift of densities defines distinct subclasses of extracellular vesicles from human whole saliva. PLoS One 16, e0249526 (2021).

23. F. Pedregosa et al., Scikit-learn: Machine learning in Python. J Mach Learn Res 12, 2825–2830 (2011).

24. C. R. Harris et al., Array programming with NumPy. Nature 585, 357–362 (2020).

25. P. Virtanen et al., SciPy 1.0: fundamental algorithms for scientific computing in Python. Nat Mthods 17, 261–272 (2020).

26. R. Tavenard et al., Tslearn, a machine learning toolkit for time series data. J Mach Learn Res 21, 1–6 (2020).

27. R. Bedre (Reneshbedre/Bioinfokit: Bioinformatics data analysis and visualization toolkit. doi:10.5281/zenodo.3841708 Zenodo (2021).

28. J. D. Hunter, Matplotlib: A 2D graphics environment. Comput Sci Eng 9, 90–95 (2007).

29. M. L. Waskom, Seaborn: statistical data visualization. J Open Source Softw 6, 3021 (2021).

30. J. Reback et al., pandas-dev/pandas: Pandas 1.0.5. doi:10.5281/zenodo.3898987 Zenodo (2020).

31. M. Ashburner et al., Gene ontology: tool for the unification of biology. The Gene Ontology Consortium. Nat Genet 25, 25–29 (2000).

32. C. Gene Ontology, The Gene Ontology resource: enriching a GOld mine. Nucleic Acids Res 49, D325–D334 (2021).

33. Y. Zhou et al., Metascape provides a biologist-oriented resource for the analysis of systems-level datasets. Nat Commun 10, 1523 (2019).

34. E.-T. Consortium et al., EV-TRACK: transparent reporting and centralizing knowledge in extracellular vesicle research. Nat Methods 14, 228–232 (2017).

35. B. Carr, P. Hole, A. Malloy, P. Nelson, J. Smith, Applications of nanoparticle tracking analysis in nanoparticle research--A mini-review. Eur J Parenter Pharm Sci 14, 45 (2009).

36. R. A. Dragovic et al., Sizing and phenotyping of cellular vesicles using nanoparticle tracking analysis. Nanomedicine 7, 780–788 (2011).

37. E. van der Pol et al., Particle size distribution of exosomes and microvesicles determined by transmission electron microscopy, flow cytometry, nanoparticle tracking analysis, and resistive pulse sensing. J Thromb Haemost 12, 1182–1192 (2014).

38. Y. Ishikawa, H. Ishida, Aquaporin water channel in salivary glands. Jpn J Pharmacol 83, 95–101 (2000).

39. I. Brody, G. Ronquist, A. Gottfries, Ultrastructural localization of the prostasome - an organelle in human seminal plasma. Ups J Med Sci 88, 63–80 (1983).

40. T. Nonaka, D. T. W. Wong, Saliva-exosomics in cancer: molecular characterization of cancer-derived exosomes in saliva. Enzymes 42, 125–151 (2017).

41. G. Valcz et al., *En bloc* release of MVB-like small extracellular vesicle clusters by colorectal carcinoma cells. J Extracell Vesicles 8, 1596668 (2019).

42. R. Linares, S. Tan, C. Gounou, N. Arraud, A. R. Brisson, High-speed centrifugation induces aggregation of extracellular vesicles. J Extracell Vesicles 4, 29509 (2015).

43. R. Xu, D. W. Greening, H. J. Zhu, N. Takahashi, R. J. Simpson, Extracellular vesicle isolation and characterization: toward clinical application. J Clin Invest 126, 1152–1162 (2016).

44. A. Ropolo et al., The pancreatitis-induced vacuole membrane protein 1 triggers autophagy in mammalian cells. J Biol Chem 282, 37124–37133 (2007).

45. P. J. Rousseeuw, Silhouettes: a graphical aid to the interpretation and validation of cluster analysis. J Comput Appl Math 20, 53–65 (1987).

46. D. K. Jeppesen et al., Reassessment of exosome composition. Cell 177, 428–445 (2019).

47. A. L. Goldberg, K. L. Rock, Proteolysis, proteasomes and antigen presentation. Nature 357, 375–379 (1992).

48. M. Bordi et al., A gene toolbox for monitoring autophagy transcription. Cell Death Dis 12, 1044 (2021).

49. C. Xu et al., Functional interaction of heat shock protein 90 and Beclin 1 modulates Toll-like receptor-mediated autophagy. FASEB J 25, 2700–2710 (2011).

50. S. J. Gould, G. Raposo, As we wait: coping with an imperfect nomenclature for extracellular vesicles. J Extracell Vesicles 2 20389 (2013).

51. A. C. Dixson, T. R. Dawson, D. Di Vizio, A. M. Weaver, Context-specific regulation of extracellular vesicle biogenesis and cargo selection. Nat Rev Mol Cell Biol 10.1038/s41580-023-00576-0 (2023).

52. C. Théry et al., Proteomic analysis of dendritic cell-derived exosomes: a secreted subcellular compartment distinct from apoptotic vesicles. J Immunol 166, 7309–7318 (2001).

53. S. Charrin et al., Differential stability of tetraspanin/tetraspanin interactions: role of palmitoylation. FEBS Lett 516, 139–144 (2002).

54. R. Umeda, T. Nishizawa, O. Nureki, Crystallization of the human tetraspanin protein CD9. Acta Crystallogr F Struct Biol Commun 75, 254–259 (2019).

55. T. Shoham et al., The tetraspanin CD81 regulates the expression of CD19 during B cell development in a postendoplasmic reticulum compartment. J Immunol 171, 4062–4072 (2003).

56. K. Ito, Identification of the *secY* (*prlA*) gene product involved in protein export in *Escherichia coli*. Mol Gen Genet 197, 204–208 (1984).

57. K. Xia et al., Identifying the subproteome of kinetically stable proteins via diagonal 2D SDS/PAGE. Proc Natl Acad Sci U S A 104, 17329–17334 (2007).

58. H. Zhang et al., Identification of distinct nanoparticles and subsets of extracellular vesicles by asymmetric flow field-flow fractionation. Nat Cell Biol 20, 332–343 (2018).

59. Q. Zhang et al., Supermeres are functional extracellular nanoparticles replete with disease biomarkers and therapeutic targets. Nat Cell Biol 23, 1240–1254 (2021).

60. K. Al-Nedawi et al., Intercellular transfer of the oncogenic receptor EGFRvIII by microvesicles derived from tumour cells. Nat Cell Biol 10, 619–624 (2008).

