## Supplemental figure for "Differentiation of Large Extracellular Vesicles in Oral Fluid: Combined Protocol of Small Force Centrifugation and Pattern Analysis"

### Figure. S1

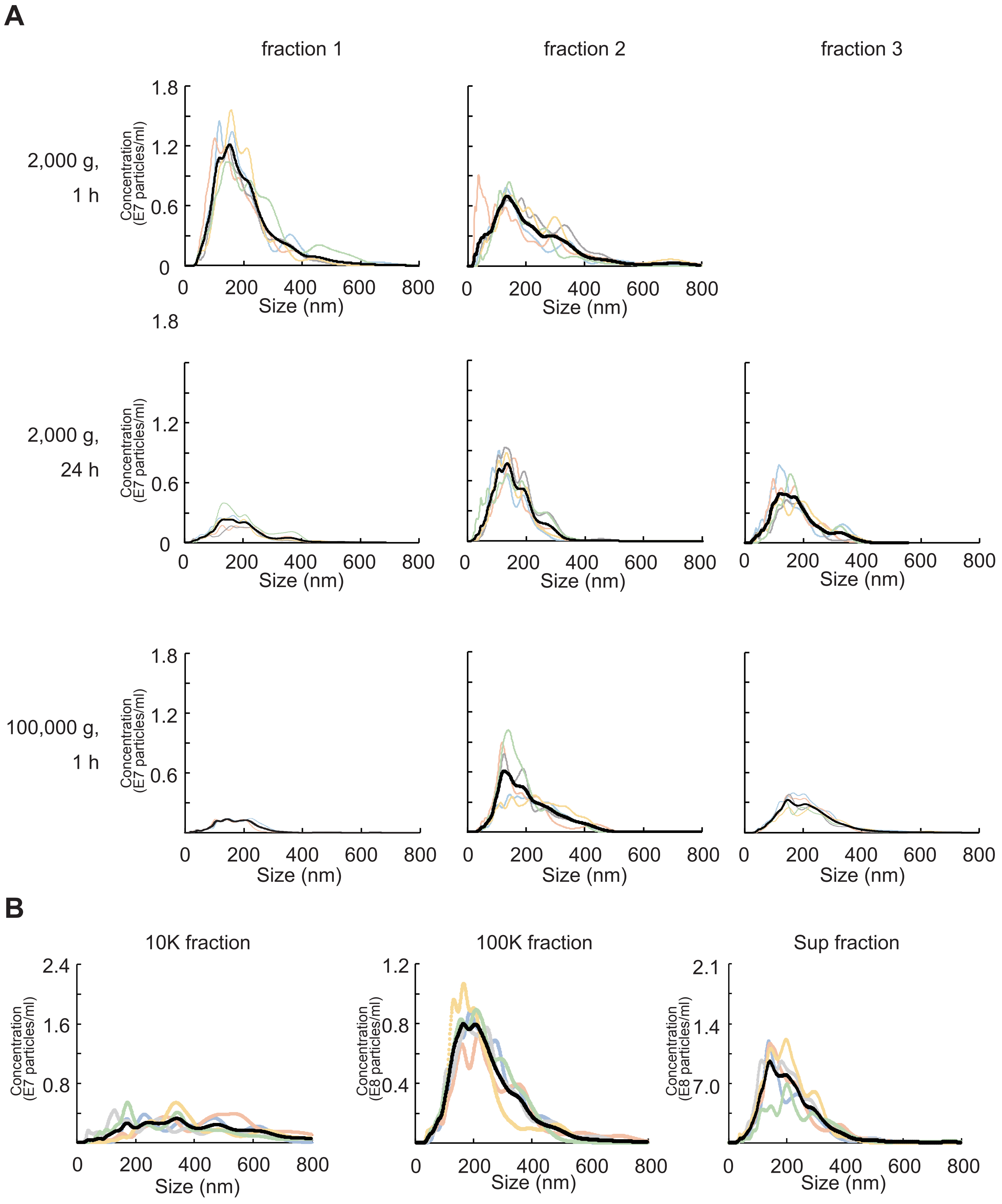
A
fraction 1
fraction 2
fraction 3
2,000 g,
1 h
2,000 g,
24 h
100,000 g,
1 h
Concentration
(E7 particles/ml)
1.8
1.2
0.6
0
0
200
400
600
800
Size (nm)
B
10K fraction
100K fraction
Sup fraction
Concentration
(E7 particles/ml)
2.4
1.6
0.8
Concentration
(E8 particles/ml)
1.2
0.8
0.4
2.1
1.4
7.0

Figure. S2

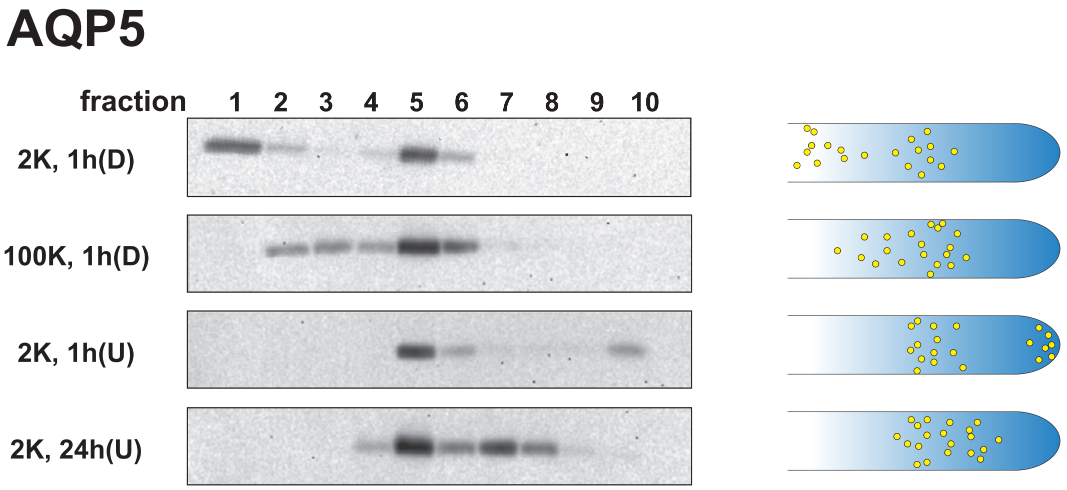

### Figure. S3

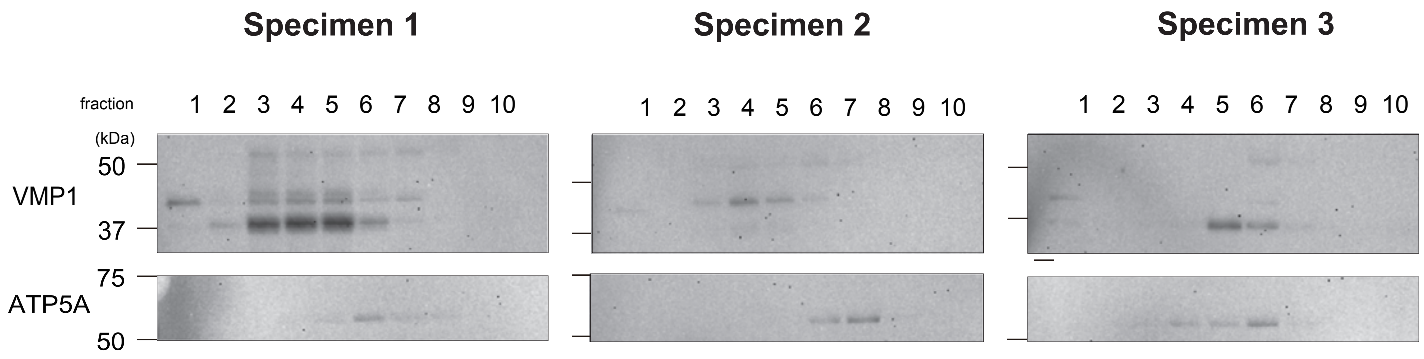
Specimen 1
Specimen 2
Specimen 3
fraction
1 2 3 4 5 6 7 8 9 10
1 2 3 4 5 6 7 8 9 10
1 2 3 4 5 6 7 8 9 10
(kDa)
50
VMP1
37
75
ATP5A
50

### Figure. S4

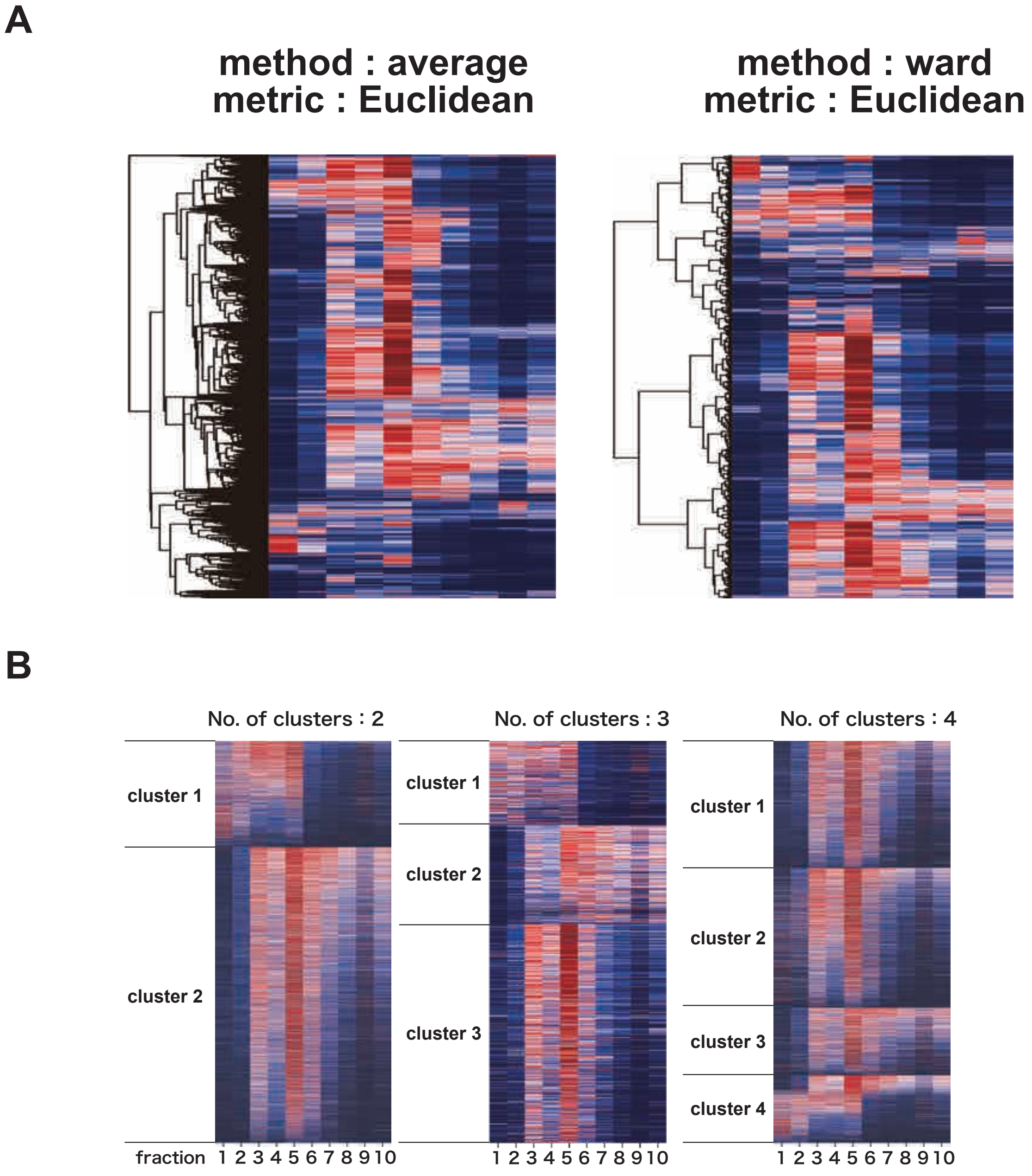
A
method : average
metric : Euclidean
method : ward
metric : Euclidean
B
No. of clusters : 2
cluster 1
cluster 2
No. of clusters : 3
cluster 1
cluster 2
cluster 3
No. of clusters : 4
cluster 1
cluster 2
cluster 3
cluster 4
fraction
1 2 3 4 5 6 7 8 9 10

### Figure. S5

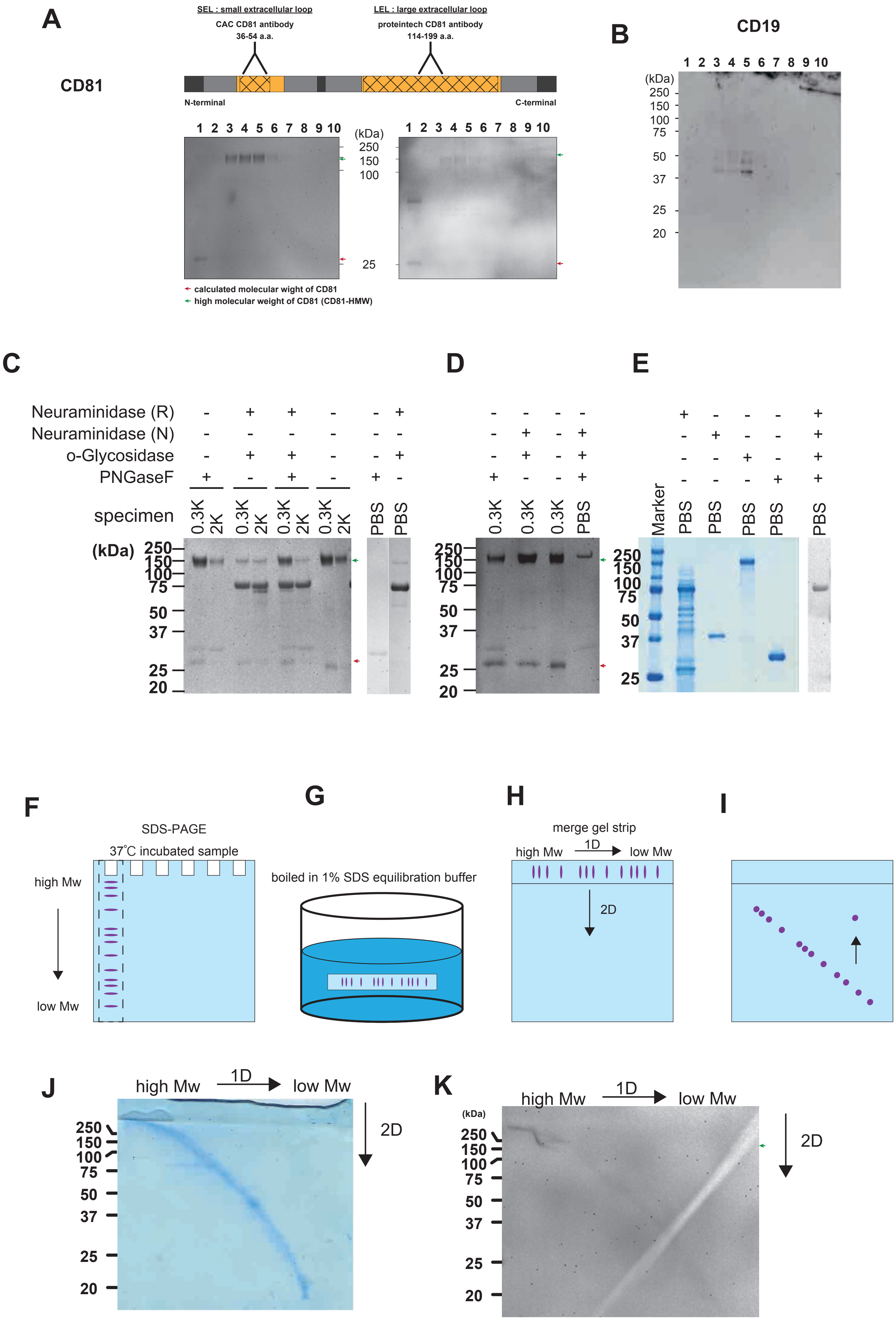
A
SEL : small extracellular loop
CAC CD81 antibody
36-54 a.a.
LEL : large extracellular loop
proteintech CD81 antibody
114-199 a.a.
CD81
N-terminal
C-terminal
1 2 3 4 5 6 7 8 9 10
(kDa)
250
150
100
25
calculated molecular wight of CD81
high molecular weight of CD81 (CD81-HMW)
B
CD19
1 2 3 4 5 6 7 8 9 10
(kDa)
250
150
100
75
50
37
25
20
C
Neuraminidase (R)
Neuraminidase (N)
o-Glycosidase
PNGaseF
specimen
0.3K 2K 0.3K 2K 0.3K 2K 0.3K 2K PBS PBS
(kDa)
250
150
100
75
50
37
25
20
D
0.3K 0.3K 0.3K PBS
E
Marker PBS PBS PBS PBS PBS
F
SDS-PAGE
37˚C incubated sample
high Mw
low Mw
G
boiled in 1% SDS equilibration buffer
H
merge gel strip
high Mw
1D
low Mw
2D
I
J
high Mw
1D
low Mw
2D
K
(kDa)

Figure. S6

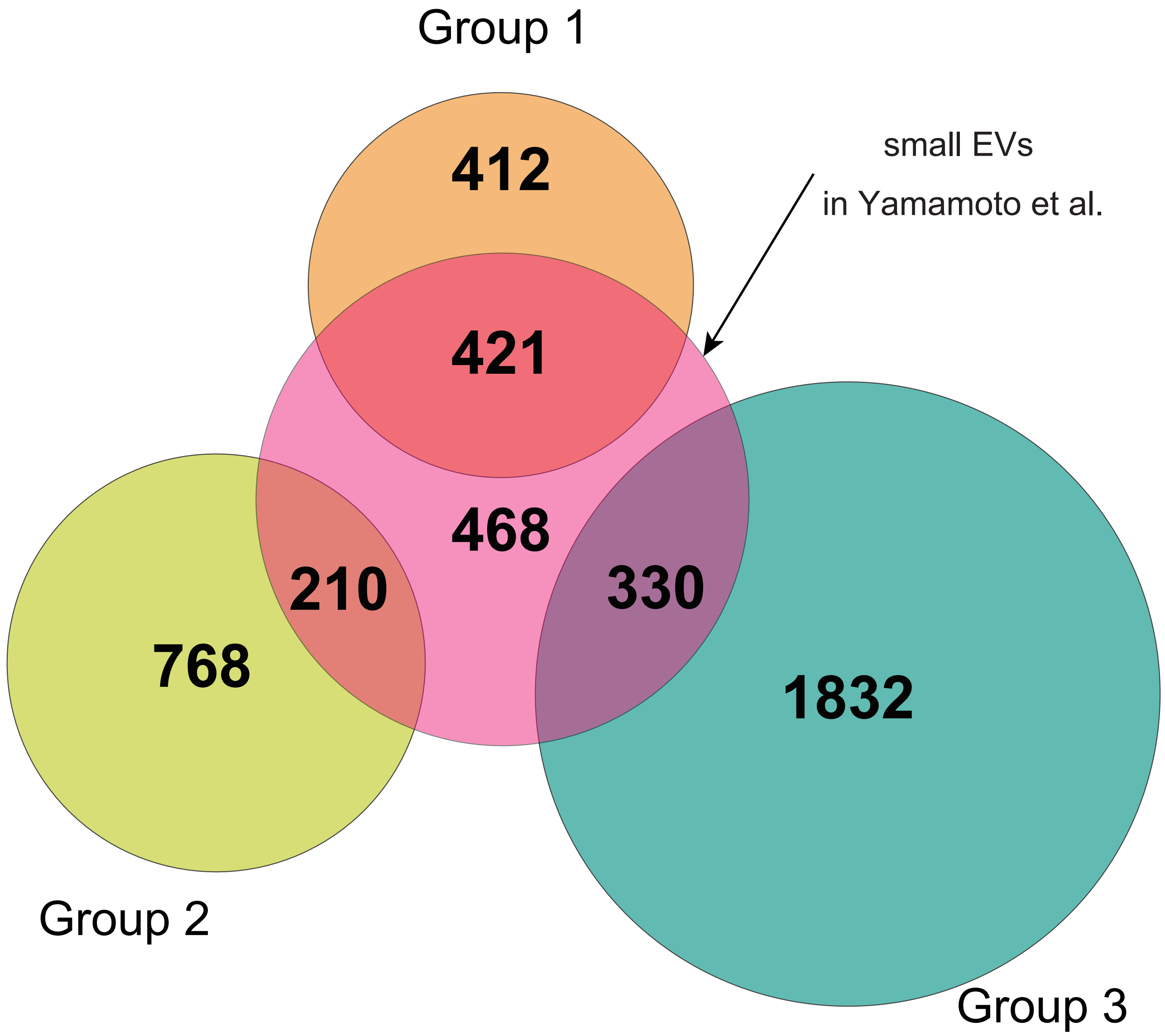
Group 1
small EVs
in Yamamoto et al.
412
421
468
210
330
768
1832
Group 2
Group 3

### Original image for western blotting

#### Fig. 2D CD9

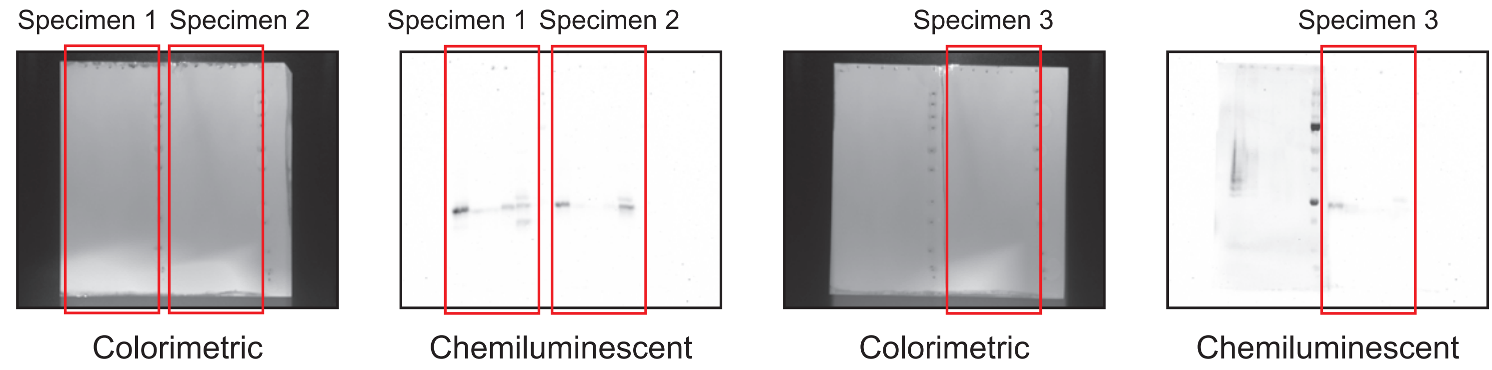


#### Fig. 2D CD63

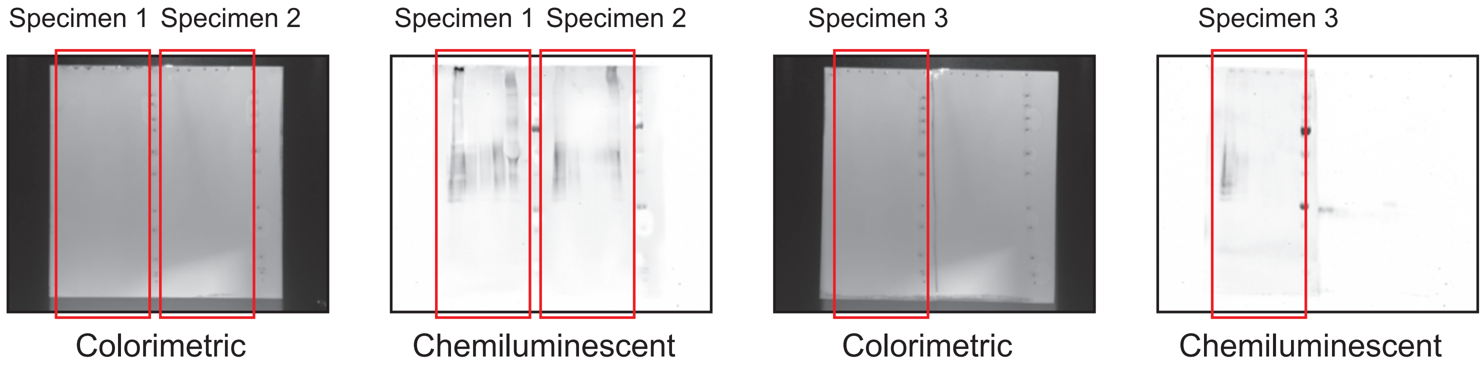


#### Fig. 2D CD81

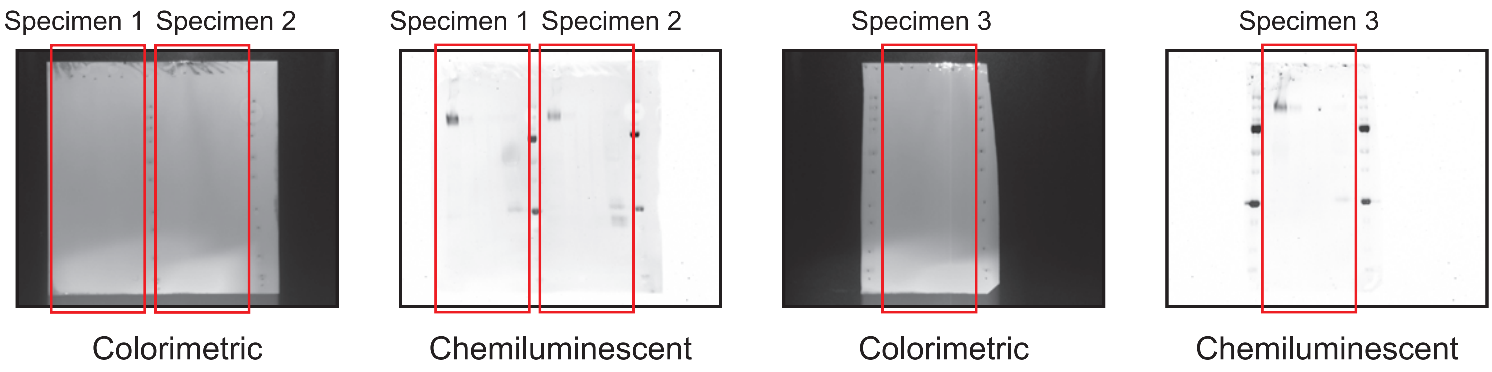

### Fig. 2E CD9

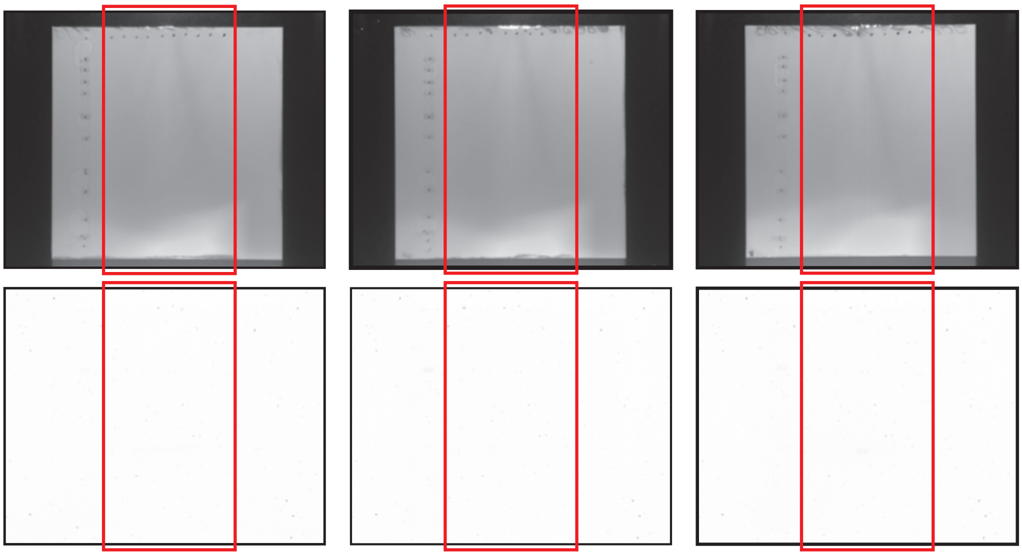


### Fig. 2E CD63

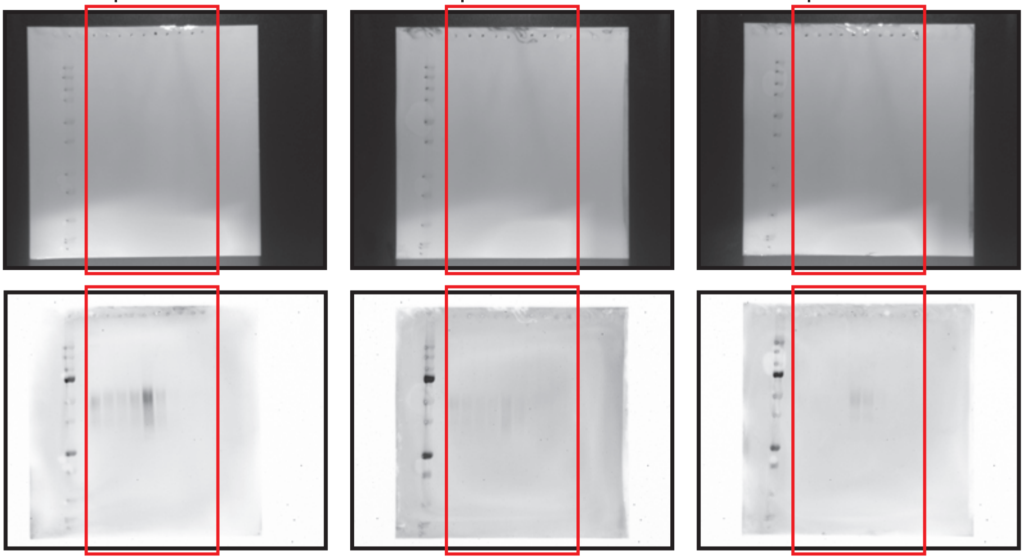


### Fig. 2E CD81

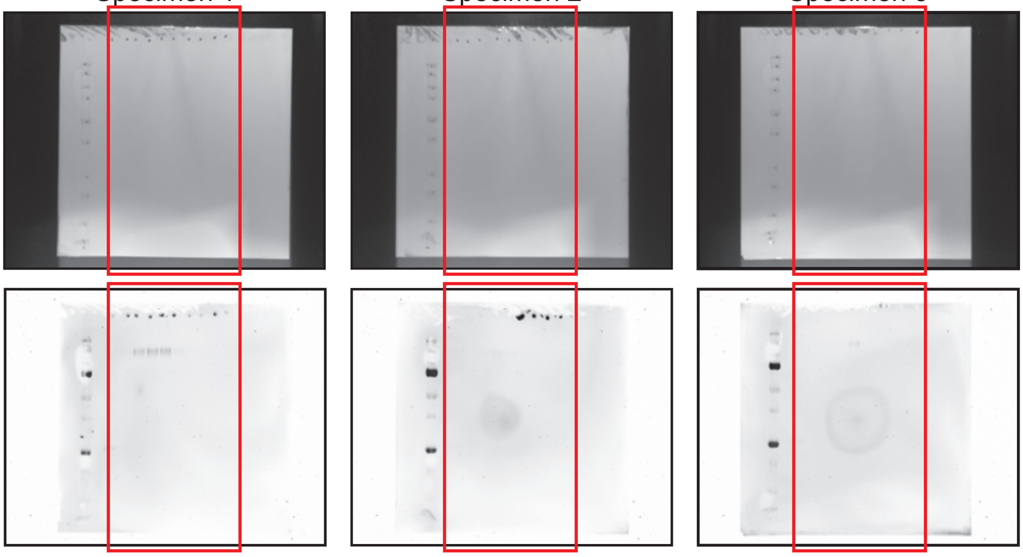

Fig. S2 AQP5

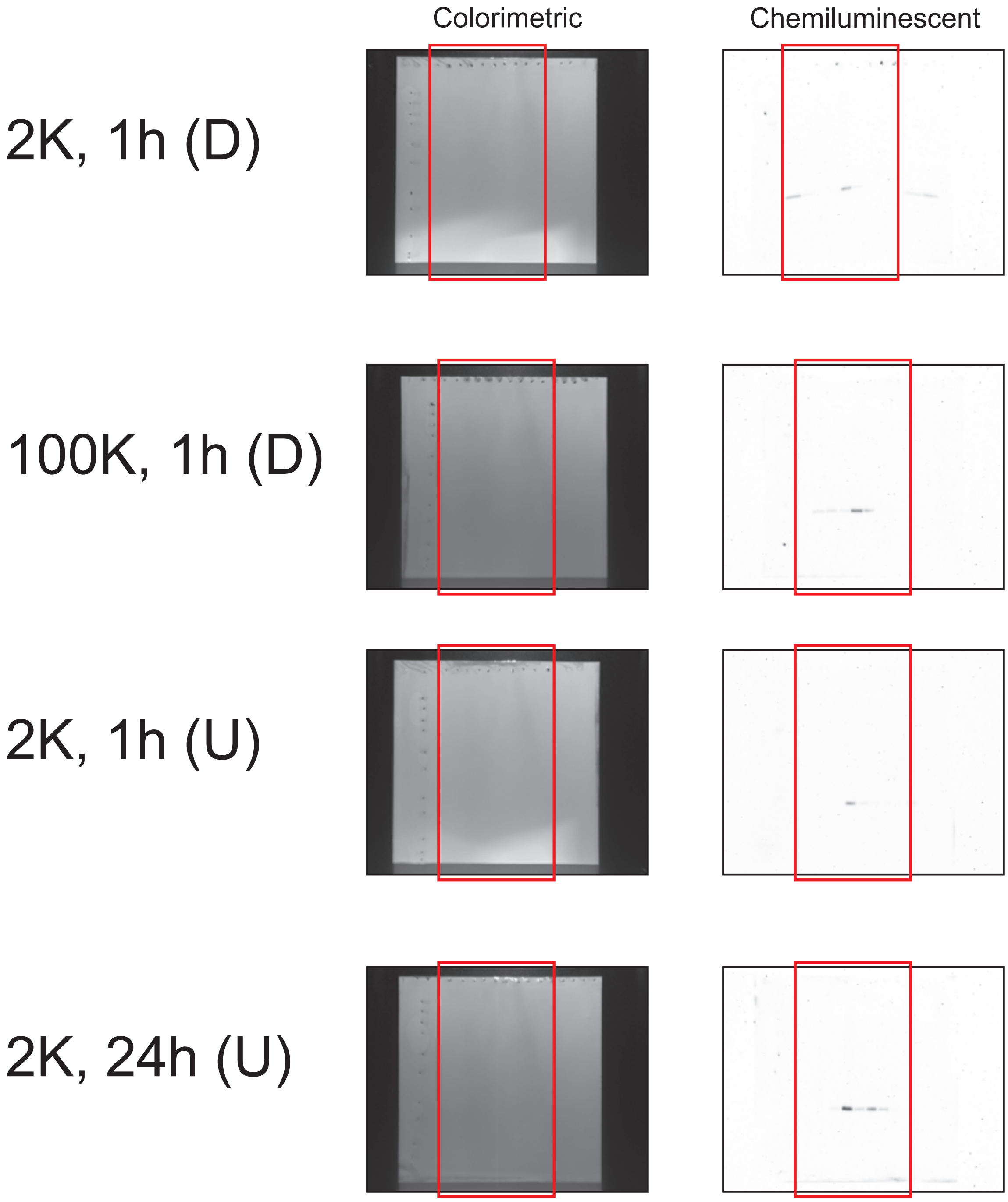

Colorimetric
Chemiluminescent
2K, 1h (D)
100K, 1h (D)
2K, 1h (U)
2K, 24h (U)

### Fig. S3 VMP1

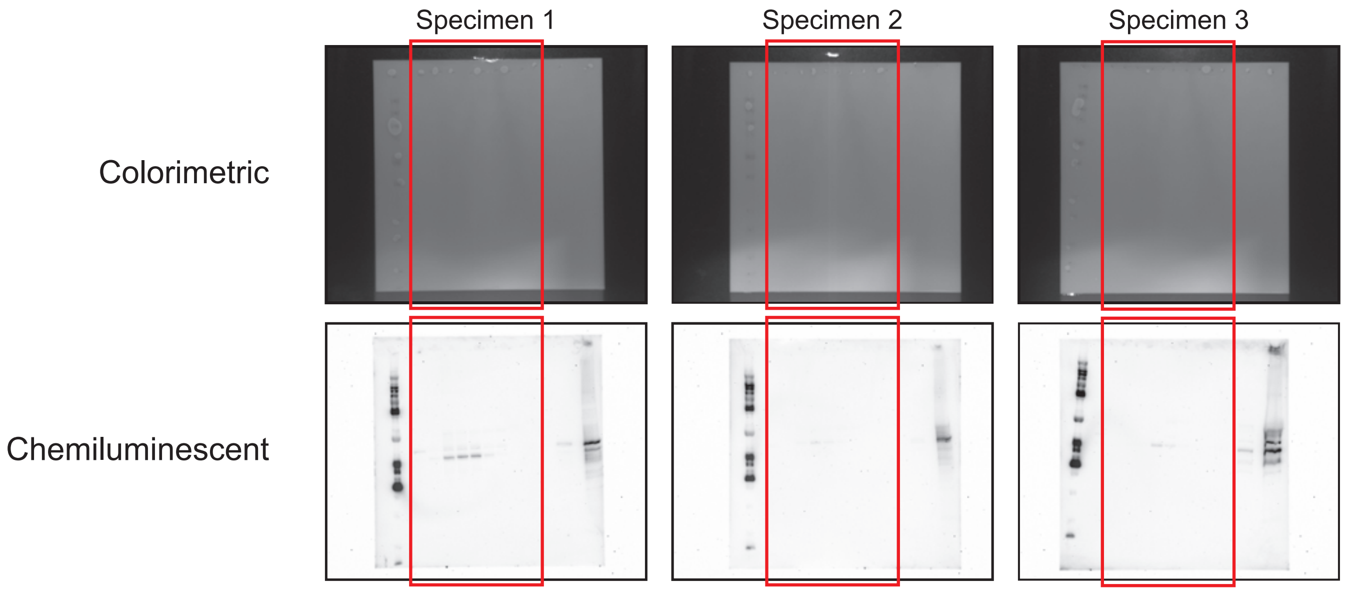


### Fig. S3 ATP5A

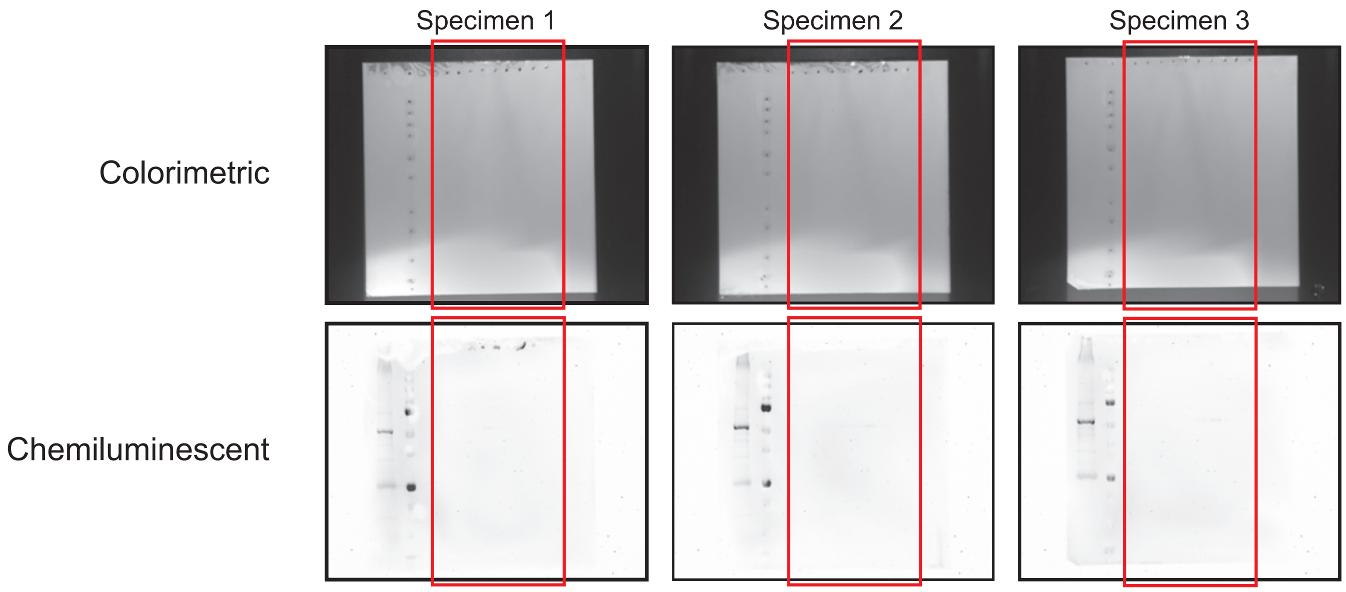

### Fig. S6A

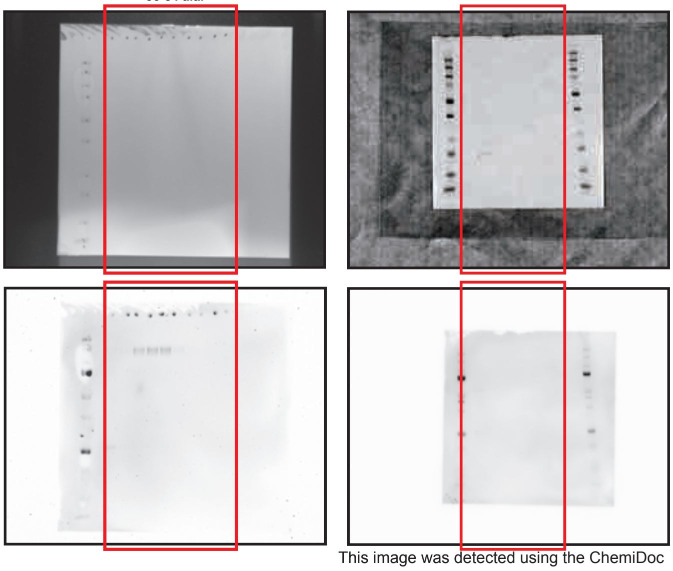


This image was detected using the ChemiDoc Touch Imaging System (17001401JA, Bio Rad).

### Fig. S6B

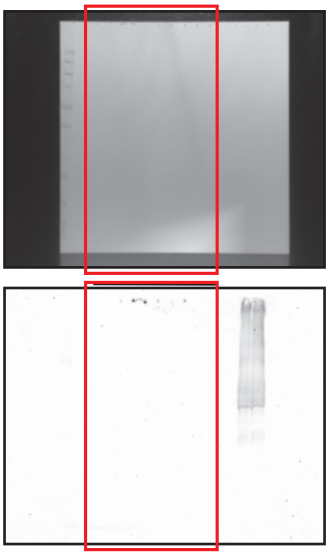

### Fig. S6C

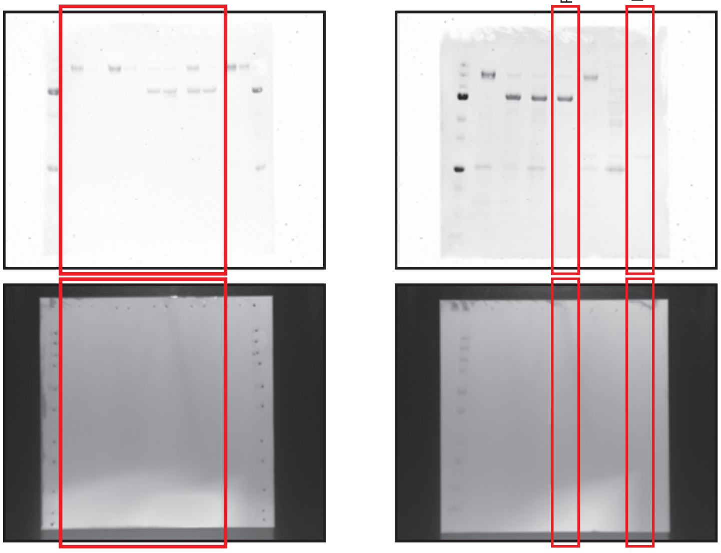


### Fig. S6D

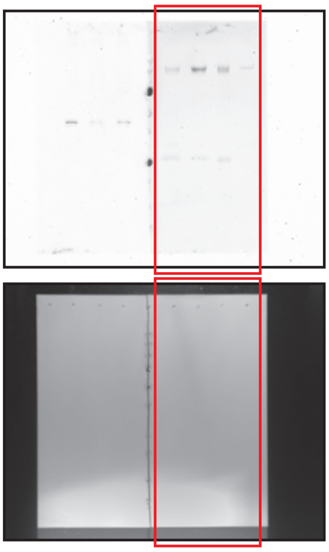


### Fig. S6E

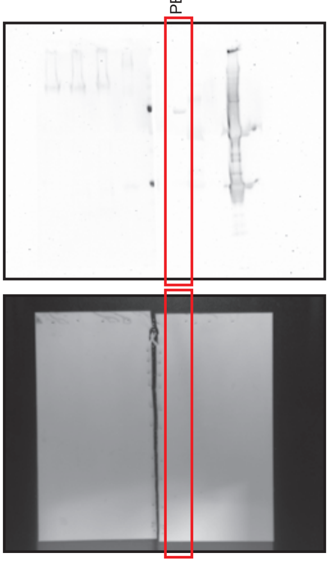


### Fig. S6K

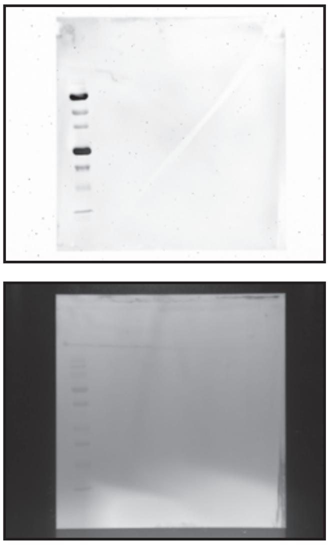
