## Supplemental information for "Differentiation of Large Extracellular Vesicles in Oral Fluid: Combined Protocol of Small Force Centrifugation and Pattern Analysis"

**Supplementary Information 1**

Stokes' law is expressed as the following equation:

$$v = \frac{d^2 \cdot (\delta - \rho)}{18\eta} \cdot r \cdot \omega^2$$

where v: sedimentation velocity of particle, d: particle diameter, δ: density of particle, ρ: density of media, η: viscosity of media, r: distance from axis of rotation, and ω: singular velocity of rotation. Stokes' law underlies the theory of centrifugation. When the media used in centrifugation is homogeneous in density, the diameter of the particles is a major factor affecting the sedimentation velocity, since it is calculated as the square of the particle diameter. This allows for the differentiation of particles mostly based on their sizes, which enables sequential differential centrifugation. However, when a density gradient medium is used, the sedimentation velocity becomes zero when the density of the particles is equal to the density of the media, which underlies the differentiation of small EV by density gradient ultracentrifugation. Note: all particles are assumed to be spherical.

**Supplementary table 1**

List of 3,973 proteins detected by mass spectrometry. From left to right, the accession number and gene name in Uniprot (1) are shown in columns A and B, respectively. From columns C to L, the expression levels of proteins in fractions 1 to 10 are shown by setting the maximum expression per fraction to 1 and the minimum to 0. In column M, the grouping determined by ESP is shown (see main text for the details).

**Supplementary table 2**

Results of the gene ontology (GO; 2, 3) enrichment analysis using all proteins in each group by TimeSeriesKMeans (4). All statistically enriched terms (GO terms) were filtered to calculate the accumulative hypergeometric p-values and enrichment. The top 20 terms of -Log10 (P-value) were used for analysis. GO enrichment analysis was performed using Metascape (5).

**Supplementary table 3**

Distributions of proteins belonging to specific protein families among Groups 1, 2, and 3. In column F, the numbers of proteins included in each group are shown.

**Figure S1. (A)** Nanoparticle tracking analysis (NTA) of concentrations and size of small particles contained in the fractions prepared by non-equilibrium density gradient centrifugation of Oral fluids (OFs) 2K crude fraction under three distinct conditions. These were the raw data for Figure 1B. Five independent measurements were performed, and their average is shown by bold lines. **(B)** Histograms of NTA after differentiation by pentapartite fractionation.

**Figure S2.** Migration patterns AQP5 after non-equilibrium density gradient centrifugation of 2K under different conditions, that is, sedimentation (downward) at 2,000 *g* (2K) for 1 h, sedimentation (downward) at 100,000 *g* (100K) for 1 h, floating (upward) at 2,000 *g* (2K) for 1 h, and floating (upward) at 2,000 *g* (2K) for 24h. On the right, schematic distribution patterns are represented by yellow filled circles (AQP5).

**Figure S3.** Migration patterns of VMP1(P072, Medical & Biological Laboratories, Tokyo, Japan; 1: 1,000) and ATP5A (ab14748, Abcam Cambridge, UK; 1: 1,000) after non-equilibrium density gradient centrifugation of 2,000 *g* (2K) for 1 h, revealed by western blotting. Numbers on the left represent molecular weights of markers (kDa). The protocol is done as described in MATERIALS AND METHODS.

**Figure S4. (A)** Results of hierarchical clustering of mass spectrometry data obtained from 10 fractions using clustering algorithm seaborn (6) with different modes, as shown. **(B)** Results of clustering using TimeSeriesKMeans by setting numbers of clustering to be 2, 3, and 4.

**Figure S5.** The top 20 protein families in each biological process, cellar component, and molecular function classification that were identified via gene ontology enrichment analyses using metascape are shown for Groups 1 (A–C), 2 (D–F), and 3 (G–I). All statistically enriched terms (GO terms) were filtered to calculate the accumulative hypergeometric p-values and enrichment factors, and the GO terms of the top 20 $-\log_{10}$(p-value) are shown. The axis of abscissas represents $-\log_{10}$(p-value), and GO terms are shown on the right.

**Figure S6. (A)** Western blot of CD81 using two antibodies that recognize different epitopes (schematically illustrated on the top). Both antibodies (SHI-EXO-M03, Cosmobio) and (66866-1-Ig,

Proteintech) detect CD81 (the apparent molecular weight is 26 kDa) and high molecular weight of CD81 (CD81-HMW) (150 K). The right western blot image is identical to the one used in Figure 2C. **(B)** Western blotting using anti-CD19 antibody (#3574, Cell Signaling Technology, MA, USA). Note: 100 K signal was invisible in this experiment. **(C–E)** Deglycosylation experiments with O-Glycosidase & Neuraminidase Bundle (E0540, NEB) and PNGaseF (P0704, NEB), Neuraminidase (269611, Roche), PNGaseF deglycosylates glycoproteins by hydrolyzing N-glycosidic bonds between asparagine and N-acetylglucosamine, whereas O-glycosidase cleaves O-glycosidic bonds between N-acetylgalactosamine and the hydroxyl group of serine or threonine. Neuraminidase was added because it is known to enhance the deglycosylation reaction by O-Glycosidase. After incubation of 0.3K and 2K fractions with the combination of these proteins, western blotting was performed with an anti-CD81 antibody (SHI-EXO-M03, Cosmobio). As shown in **C** left panel, the first O-Glycosidase seemed to decrease the molecular weight of CD81-HMW from 150K to 75K. However, this 75K signal was detected even without 0.3K and 2K if O-glycosidase was present (right panel). In experiment (**C**), we used the O-Glycosidase & Neuraminidase Bundle (E0540, NEB). Therefore, there was a possibility that neuraminidase in this mixture had the epitope for anti-CD81 antibody. Indeed, when we used neuraminidase (269611, Roche; and single O-Glycosidase, P0002, NEB), this 75K signal disappeared. Neuraminidase N and R are produced by NEB and Roche. When these enzymes were separated by SDS-PAGE and stained with Coomassie brilliant blue (CBB), neuraminidase R contained major proteins whose apparent molecular weights were 75K, indicating neuraminidase R has accidently processed the epitope that reacted with anti-CD81 antibody. After replacing the sample with PBS and probing only the secondary antibody without incubating the primary antibody, a 75-kDa signal was detected. Thus, Neuraminidase R may contain a factor that reacts with mouse IgG. **(F-I)** The effect of heat treatment on the mobility of CD81 on SDS-PAGE. In Figures F to H, the scheme of D2D-SDS-PAGE (7) is shown. Briefly, protein samples were treated with low temperature (37°C), and the first 1D separation was performed as described in Material and Methods in main text (**F**). The 1D gel was then cut into a strip, which was boiled for 10 min in 1% SDS equilibration buffer (50 mM Tris-HCl, 1% SDS, 15% glycerol, 0.02% bromophenol blue, pH 6.8, 100°C) (**G**). The strip of (G) was placed on the top of an 80 mm × 80 mm × 1 mm gel of 12.5 (w/v) % [prepared with acrylamide/bis mixed solution (29:1) (06141-35, Nacalai), 375 mM Tris-HCl (pH 8.8), 0.1% SDS, 0.033% Ammonium peroxodisulfate (APS, 02602-02, Nacalai), and 0.05% N,N,N',N'-Tetramethylethylenediamine (TEMED, 33401-72, Nacalai)] having a 4.75 (w/v) % stacking gel [prepared with acrylamide/bis mixed solution (29:1), 125mM Tris-HCl (pH 6.8), 0.1% SDS, 0.033 % APS, 0.1% TEMED], which was prepared using a glass plate set (01-210, TEFCO, Tokyo, Japan) and electrophoresed using an electrophoresis chamber, XCell SureLock (EI0002, invitrogen) (**H**). Proteins that migrated diagonally were not affected by heat treatment, whereas ones that deviated from this diagonal line must have apparent molecular weights that are affected by the boiling step in (G) (**I**). After the gel was stained with CBB (J), separated proteins were transferred to the membrane and probed with anti-CD81 antibody (SHI-EXO-M03, Cosmobio), as described in Material and Methods (main text) (**K**). In this D2D-PAGE experiment, only the 150K signal was observed around the diagonal line and no signal appeared around 25K in the second SDS-PAGE **(K)**, suggesting the CD81-HMW moiety was not an artifact derived from the boiling treatment. Red and green arrows indicate the calculated molecular weight of CD81 and the high molecular weight of CD81 (CD81-HMW), respectively.

### SUPPLEMENTARY REFERENCES


1. C. UniProt, UniProt: the Universal Protein Knowledgebase in 2023. *Nucleic Acids Res* **51**, D523-D531 (2023).
2. C. Gene Ontology, The Gene Ontology resource: enriching a GOld mine. *Nucleic Acids Res* **49**, D325-D334 (2021).
3. M. Ashburner *et al.*, Gene ontology: tool for the unification of biology. The Gene Ontology Consortium. *Nat Genet* **25**, 25-29 (2000).
4. R. Tavenard *et al.*, Tslearn, a machine learning toolkit for time series data. *J Mach Learn Res* **21**, 1-6 (2020).
5. Y. Zhou *et al.*, Metascape provides a biologist-oriented resource for the analysis of systems-level datasets. *Nat Commun* **10**, 1523 (2019).
6. M. L. Waskom, Seaborn: statistical data visualization. *J Open Source Software* **6**, 3021 (2021).
7. K. Xia *et al.*, Identifying the subproteome of kinetically stable proteins via diagonal 2D SDS/PAGE. *Proc Natl Acad Sci U S A* **104**, 17329-17334 (2007).
